# Genetically variant human pluripotent stem cells selectively eliminate wild-type counterparts through YAP-mediated cell competition

**DOI:** 10.1101/854430

**Authors:** Christopher J. Price, Dylan Stavish, Paul J. Gokhale, Samantha Sargeant, Joanne Lacey, Tristan A. Rodriguez, Ivana Barbaric

## Abstract

The appearance of genetic changes in human pluripotent stem cells (hPSCs) presents a concern for their use in research and regenerative medicine. Variant hPSCs harbouring recurrent culture-acquired aneuploidies display growth advantages over wild-type diploid cells, but the mechanisms yielding a drift from predominantly wild-type to variant cell populations remain poorly understood. Here we show that the dominance of variant clones in mosaic cultures is enhanced through competitive interactions resulting in elimination of wild-type cells. This elimination occurs through corralling and mechanical compression by faster growing variants, causing a redistribution of F-actin and sequestration of YAP in the cytoplasm that induces apoptosis in wild-type cells. Importantly, YAP overexpression in wild-type cells is sufficient to alleviate their loser phenotype. Our results demonstrate that hPSC fate is coupled to mechanical cues imposed by neighbouring cells and reveal that hijacking this mechanism allows variants to achieve clonal dominance in cultures.

## Introduction

Cell-cell interaction is a critical feature of multicellular organisms, necessary for the orderly development of tissues and maintenance of their homeostasis. The ability of cells to influence their neighbouring cells’ fate choices has become apparent from studies in various *in vitro* and *in vivo* models. An example of this is cell competition, a type of cell-cell interaction wherein viable but less-fit “loser” cells are outcompeted for nutrients or space and eventually eliminated by the fitter “winner” cells (reviewed in (Bowling et al., 2019)). Initially described and studied in *Drosophila* as a tissue homeostatic mechanism (Morata and Ripoll, 1975), over recent years it has become evident that a form of cell competition, known as super-competition, is implicated in expansion of cancerous cells (Eichenlaub et al., 2016; Suijkerbuijk et al., 2016). In super-competition, the acquisition of a mutation which enhances the relative fitness of a cell results in the removal of neighbouring wild-type cells (Johnston, 2014).

In the context of regenerative medicine, the fundamental question of how mutant cells may influence behaviour of their wild-type counterparts has been brought into focus by observation that human pluripotent stem cells (hPSCs) acquire genetic changes upon prolonged passaging (Draper et al., 2004; International Stem Cell et al., 2011). Studies of genetic integrity of hPSCs over the last two decades have revealed a bias in genetic changes acquired in hPSCs, with the most common karyotypic abnormalities involving gains of chromosomes 1, 12, 17, 20 and X (Baker et al., 2007; Draper et al., 2004; International Stem Cell et al., 2011). The recurrent nature of genetic abnormalities in hPSCs is indicative of such changes conferring selective growth advantage to the variant cells (Baker et al., 2007; Draper et al., 2004). The implications of the variant presence could be significant for therapeutic and research uses of hPSCs, as altered behaviour of variant cells could impact on the efficiency of differentiation protocols, functionality of differentiated cells or the safety of cell replacement therapies (Andrews et al., 2017). Of particular safety concern is the observation that aneuploidies commonly observed in hPSCs, such as the gain of chromosomes 12 or 17, are also characteristic of the malignant PSCs of germ cell tumours, teratocarcinomas (Andrews et al., 2005; Harrison et al., 2007). Hence, resolving the mechanisms that lead to genetic changes and their subsequent overtake of hPSC culture is pivotal for informing approaches to minimise the appearance of genetic variants in culture.

The emergence of variant cells in hPSC cultures has been likened to the process of evolution, whereby the interplay of mutation and selection leads to the expansion of clones which possess the greatest growth advantage under particular culture conditions (Andrews et al., 2005). Indeed, selective advantage of commonly occurring genetic changes in hPSCs has been demonstrated through mixing experiments, wherein spiking a small proportion of variant cells into wild-type cultures resulted in a rapid overtake of cultures by the variants (Avery et al., 2013; Olariu et al., 2010). To explain the reasons behind the variant overtake of cultures, studies of variant cells have mostly focused on the intrinsic properties that could lead to their growth advantage, such as enhanced proliferation and reduced levels of apoptosis (Avery et al., 2013; Barbaric et al., 2014; Ben-David et al., 2014; Draper et al., 2004; Enver et al., 2005; Nguyen et al., 2014). Yet, when the variant cells first emerge, they co-exist within the same culture as the wild-type cells, and hence share the culture environment as well as a proportion of their cell-cell contacts. However, little is known about the nature of cell-cell interactions of wild-type and variant cells in mixed cultures and whether the presence of variants affects the growth and survival of wild-type hPSCs.

Here, we show that an important aspect of the competitive advantage displayed by some of the commonly occurring variant hPSCs is the ability to induce apoptosis of wild-type cells in mosaic cultures, akin to the super-competition-like behaviour described in other cell types (de la Cova et al., 2004; Moreno and Basler, 2004). The elimination of loser cells in hPSC cultures is exerted through mechanical cues, and is mediated by YAP, downstream of the actomyosin cytoskeleton. Our findings illuminate the reliance of hPSC fates on their mechanical environment and highlight the need for consideration of culture space limitations in the scale up of hPSCs for research or clinical use.

## Results

### Variant hPSCs selectively eliminate diploid wild-type counterparts from co-cultures

To uncover the reasons behind the rapid overtake of cultures by genetically variant hPSCs (Olariu et al., 2010), we sought to examine how wild-type and genetically variant hPSCs interact and whether they affect each other’s growth. To this end, we initially used two diploid H7 sublines (either non-modified or genetically engineered to constitutively express red fluorescent protein (RFP), termed wild-type and wild-type-RFP, respectively), and their aneuploid variant harbouring a gain of chromosomes 1, 12, 17q and 20q CNV, and stably expressing green fluorescent protein (GFP) (termed variant-GFP). Time-lapse microscopy of co-cultures containing wild-type-RFP and variant-GFP cells showed a selective elimination of wild-type-RFP cells during a three-day culture period (**Video S1**). To establish that the observed elimination is due to the presence of variant cells in mixed cultures we compared the growth rates of wild-type-RFP or unlabelled wild-type cells in separate culture to how they grew in mixed cultures with variant-GFP cells. Wild-type sublines were viable and created well-established, large colonies in separate culture, but consistent with previous findings, did grow slower than variant cells (**Figure 1A**; Figure S1A) (Barbaric et al., 2014; Enver et al., 2005). In contrast to this, strikingly, upon mixing the equal numbers of variant-GFP cells, the wild-type-RFP or unlabelled wild-type cells showed severely compromised growth (**Figure 1B-D**; Figure S1B-D). The co-culture had no effect on the number of variant-GFP cells (**Figure 1A,B**; Figure S1A,B). We confirmed that the same competitive interaction occurred in another pair of diploid and aneuploid cells of a different hPSC line (H14), with aneuploid cells also outcompeting diploid cells in co-cultures (**Figure S2**). Overall, these experiments demonstrated that the presence of variant cells negatively affects the numbers of wild-type cells in co-cultures with variants.

**Figure 1.**
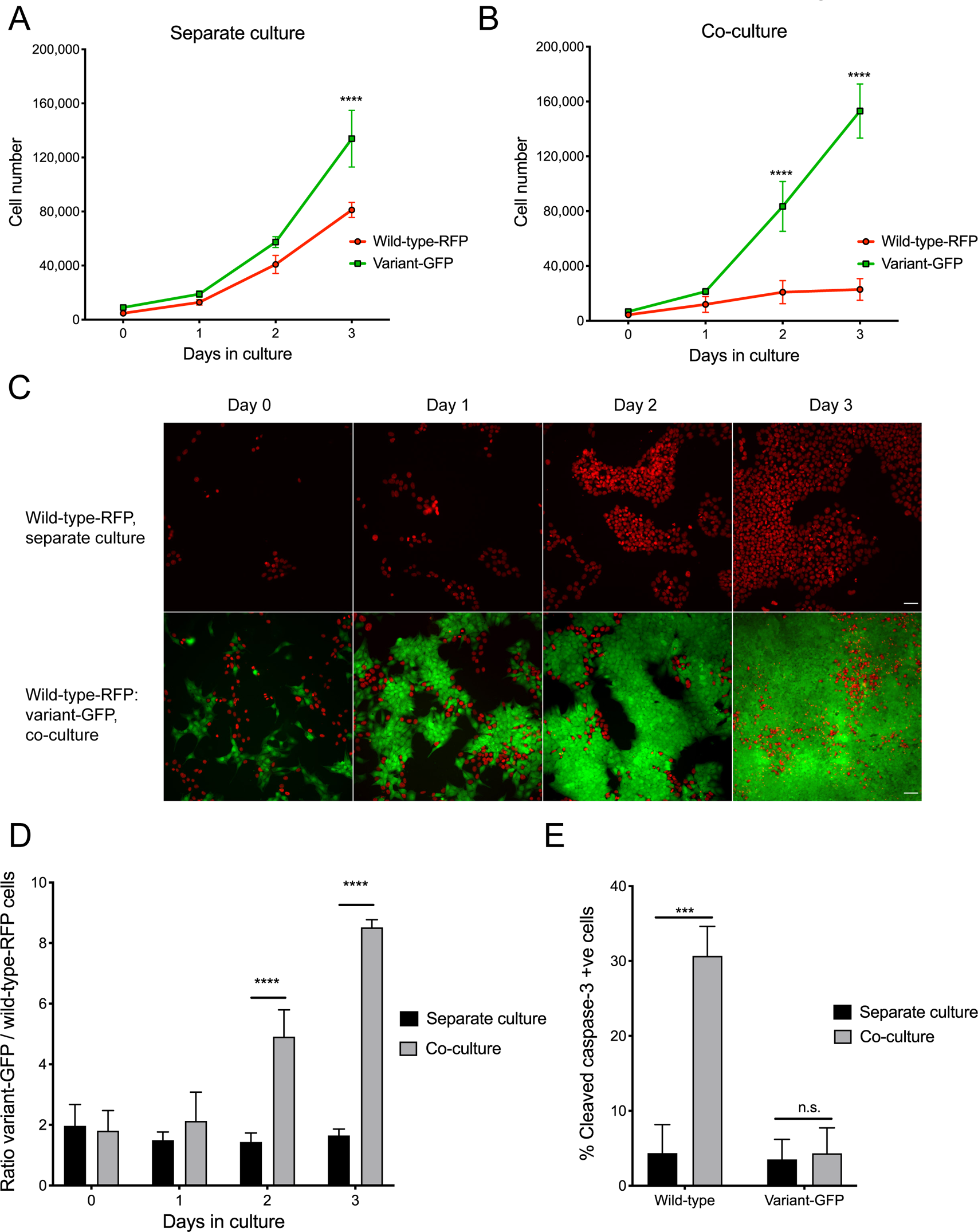
Wild-type cells are eliminated by apoptosis from co-cultures with variant hPSCs. A) Growth curves of wild-type-RFP and variant-GFP cells grown separately. B) Growth curves of wild-type-RFP and variant-GFP cells grown in co-culture. C) Representative images of wild-type-RFP hPSCs (red) grown separately (upper panels) or in co-culture with variant-GFP hPSCs (green) (lower panels). Scale bar: 50μm. D) Ratio of variant-GFP/wild-type-RFP cells in separate versus co-culture conditions. ‘Separate culture’ ratio was calculated by dividing the number of variant-GFP cells in separate culture with a number of wild-type-RFP cells in separate culture. ‘Co-culture’ ratio was obtained by directly counting the number of either wild-type-RFP or variant-GFP cells in co-culture using high-content microscopy and dividing by the total cell count. E) Percentage of cells positive for cleaved caspase-3 indicator of apoptosis in wild-type and variant-GFP cells in separate culture or upon co-culture. Data represents the mean of three independent experiments ± SD. n.s. non-significant; *** p<0.001; **** p<0.0001, Student’s *t* test.

The elimination of wild-type cells which occurred in co-culture with variant cells is reminiscent of cell competition described in many different systems, whereby ‘weaker’ *loser* cells are eliminated in the presence of ‘fitter’ *winner* cells (reviewed in (Bowling et al., 2019)). Cell competition typically involves inducing either senescence (Bondar and Medzhitov, 2010) or apoptosis (Brumby and Richardson, 2003; Moreno et al., 2002; Sancho et al., 2013) in loser cells. From our time-lapse analysis of wild-type-RFP cells co-cultured with either variant-GFP cells or unlabelled wild-type cells as a control, it was evident that the loser cells were not arresting in co-cultures (**Figure S3**; **Figure S4**). On the other hand, using cleaved caspase-3 staining as a readout of apoptosis, we observed that whilst wild-type and variant cells showed similar levels of cell death in separate culture, the proportion of apoptotic cells was significantly increased in wild-type cells upon co-culture with variants (**Figure 1E**). There was no change in the cleaved caspase-3 levels of variant-GFP cells upon co-culture with wild-type hPSCs (**Figure 1E**). Based on these results, we concluded that the presence of variant cells is inducing apoptosis and thereby elimination of wild-type cells from mosaic cultures.

### Crowding of loser cells within mosaic cultures induces loser cell apoptosis

Given the apparent selective elimination of wild-type cells when co-cultured with variant-GFP hPSCs, we wanted to establish whether the increased death rate of wild-type cells was mediated through cell-cell contacts or by cell-secreted diffusible factors, or the combination of both. To address these possibilities, we first made use of a Transwell assay (Boyden, 1962) to spatially separate the two populations whilst allowing the free exchange of secreted factors in the culture media. In these conditions, the presence of variant-GFP cells did not increase the levels of activated caspase-3 staining in wild-type cells compared to when wild-type cells were co-cultured with wild-type cells in Transwell cultures (**Figure 2A**), indicating that the effect of variants on wild-type cells is not mediated by soluble factors. In contrast to this, plating the increasing ratios of variant-GFP cells in co-cultures (from 10% to 90%) in a monolayer caused an increasing suppression of wild-type cell numbers (**Figure 2B**). This suppression of the wild-type cells’ growth was density-dependent (**Figure 2C**) and was accompanied by increased cleaved caspase-3 staining (**Figure 2C**; Figure S5A). Together, these results demonstrate that cell competition in hPSC cultures is mediated by cell contact rather than by soluble factors.

**Figure 2.**
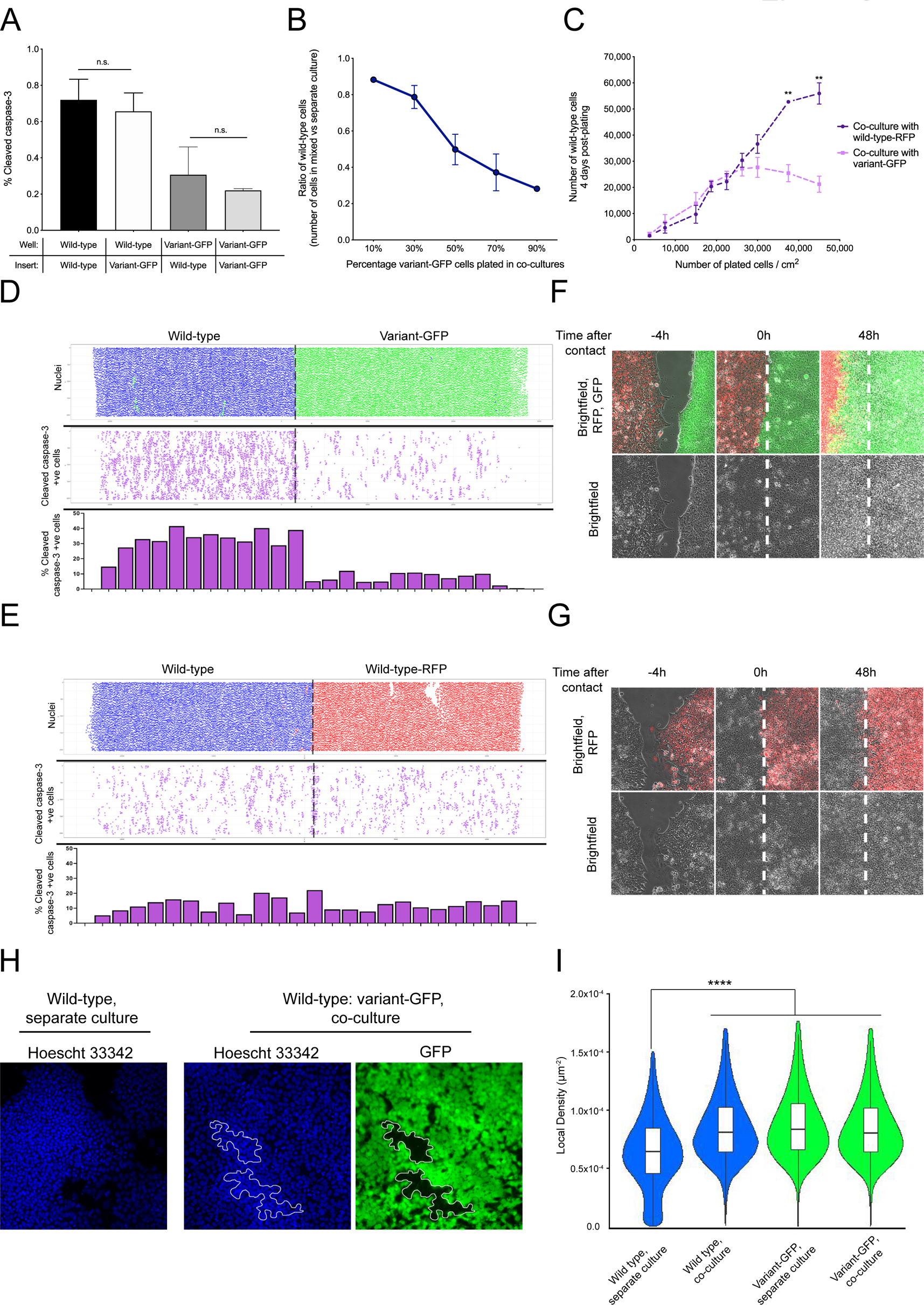
Cell competition in hPSC cultures is cell-contact mediated. A) Percentage of caspase-3 positive cells upon Transwell cultures of different sublines. Results are the mean of three independent experiments ± SD; n.s. non-significant, Student’s *t* test. B) Effect of increasing the ratio of variant-GFP cells in co-cultures with wild-type cells on the numbers of wild-type cells at day 3 of cell competition assay. Results are the mean of three independent experiments ± SD. C) Effect of increasing plating cell density on the numbers of wild-type cells. Results are the mean of three independent experiments ± SD. ** p<0.01, Student’s *t* test. D) Cell confrontation assay of wild-type-RFP and variant-GFP cells at 48h post-contact. Top panel: nuclei of wild-type-RFP and variant-GFP cells represented as blue and green dots, respectively. Middle panel: cleaved caspase-3 positive cells represented as purple dots. Bottom panel: percentage of cleaved caspase-3 positive cells calculated as the number of cleaved caspase 3-positive cells in the total cell number within a defined area of a cell insert. The width of the bar corresponds to the analysed area of the insert shown in the middle panel above. E) Cell confrontation assay of wild-type and wild-type-RFP cells at 48h post-contact. Top panel: nuclei of wild-type and wild-type-RFP cells represented as blue and red dots, respectively. Middle panel: cleaved caspase-3 signal, represented as purple dots. Bottom panel: percentage of cleaved caspase-3 positive cells calculated as the number of cleaved caspase 3-positive cells in the total cell number within a defined area of a cell insert. The width of the bar corresponds to the analysed area of the insert in the middle panel above. F) Frozen frames from the time-lapse videos of cell confrontation assay of wild-type-RFP (red) and variant-GFP (green) cells. Left panel: inserts at 4h before contact; middle panel: inserts at the time when cells first come into contact (denoted as 0h); right panel: inserts at 48h post-contact. Dashed white line indicates the position on the insert where the two different populations first meet at 0h time point. G) Frozen frames from the time-lapse videos of cell confrontation assay of wild-type and wild-type-RFP (red) cells. Left panel: inserts at 4h before contact; middle panel: inserts at the time when cells first come into contact (denoted as 0h); right panel: inserts at 48h post-contact. Dashed white line indicates the position on the insert where the two different populations first meet at 0h time point. H) Corralling of wild-type cells by variant-GFP counterparts. The outlined areas in the middle and right panels indicate regions of co-culture harbouring wild-type cells. Nuclei are counterstained with Hoechst 33342. I) Cell density of wild-type and variant-GFP cells grown either separately or in co-cultures calculated by using the nuclei of each cell as individual points to construct a Delaunay triangulation. Local density was calculated by summing the areas of Delaunay triangles sharing a vertex with the cell of interest, then taking the inverse of this sum. Data are the values of individual cells from 3 independent experiments. **** p<0.0001, one-way ANOVA.

The increased loss of wild-type cells which we observed upon increasing the ratio of variant cells in co-cultures, or upon plating the co-cultures at increasing cell densities, could be explained by two possibilities: either the higher numbers of wild-type-variant heterotypic cell contacts result in receptor-mediated cell competition (Burke and Basler, 1996), or alternatively, the winner cells are mechanically compressing the losers causing their eradication from cultures in a process termed mechanical cell competition (Levayer et al., 2016; Wagstaff et al., 2016). To distinguish between these possibilities, we first performed a cell confrontation assay, which allows two cell populations to be brought into contact at a clearly defined border (Moitrier et al., 2019; Porazinski et al., 2016). We reasoned that the receptor-mediated competition would result in cell apoptosis localised at the border of heterotypic cell contacts, whereas mechanical cell competition would result in the apoptotic signal spread throughout the areas of cell crowding (Bras-Pereira and Moreno, 2018). We plated wild-type-RFP and variant-GFP cells within separate chambers of a commercially available culture insert and allowed them to populate the area within their respective chambers overnight. Upon removal of the insert, the cells from different chambers were allowed to come into contact with each other and were then cultured for a further 48h, prior to fixing and staining for the apoptotic marker cleaved caspase-3. Supporting the notion that cell competition in hPSC cultures is mediated through mechanical means, we found that the cleaved caspase-3 was distributed within the wild-type-RFP cells beyond the heterotypic border with variant-GFP cells (**Figure 2D**). This effect was specifically caused by the presence of variants in the cell confrontation assay, as inserts containing wild-type-RFP cells and unlabelled wild-type counterparts resulted in less caspase-3 staining compared with wild-type-RFP: variant-GFP confrontation cultures (**Figure 2E**). Moreover, time-lapse imaging of the cell fronts from the time of contact over the subsequent 48h also revealed that the wild-type-RFP hPSCs were pushed back by the advancing variant-GFP population (**Figure 2F**; **Video S2**), whereas the wild-type: wild-type-RFP cells boundary remained in a similar position over 48h of tracking (**Figure 2G**; **Video S3**). Together, these results suggest that the variant-GFP cells are mechanically superior to wild-type cells and outcompete them in the competition for space.

The competition for space that we detected in the cell confrontation assays was also evident in mosaic co-cultures grown in a monolayer, as tracking of wild-type-RFP and variant-GFP cells by time lapse microscopy uncovered an apparent corralling and subsequent elimination of wild-type cells by the faster growing variants (**Video S4)**. We confirmed this effect by analysing the relative cell density of wild-type and variant-GFP cells in separate cultures and upon co-culture. The nuclei of wild-type cells in co-cultures clustered within areas of increased local density compared to the density within the separate wild-type culture (**Figure 2H,I**) and underwent apoptosis, as indicated by increased cleaved caspase-3 positive staining of corralled wild-type cells (Figure S5B). Together, these results suggest that the corralling of wild-type cells by mechanically stronger variants within mosaic cultures causes them to crowd into areas of high local cell density and thereafter commit to apoptosis.

### Winner status is conferred onto cells by having a relatively higher proliferative ability

Given that a key feature of mechanical cell competition is the crowding of loser cells caused by the faster growing winners (Levayer et al., 2016; Wagstaff et al., 2016), we next asked whether the winner status in hPSC cultures is conferred onto cells by the ability to expand faster and thus fill the available space. To this end, we performed mixing experiments of H7 wild-type-RFP cells with a range of H7 variant sublines, which harboured distinct genetic changes and displayed diverse growth rates. For example, variant H7 sublines with a gain of 1q (from herein *v1q*) or a gain of 20q copy number variant (from herein *v20q*) had similar growth rates to wild-type-RFP hPSCs in separate cultures (**Figure 3A**; Figure S6A). As predicted, upon mixing with wild-type cells, the numbers of wild-type or variant cells (either *v1q* or *v20q*) remained unaffected (**Figure 3B**; Figure S6B). In addition, we tested the behaviour of another variant line (harbouring gains of chromosome 1, 17q and isochromosome 20q (termed *v1,17q,i20*)) in co-culture with variant-GFP cells, as both lines grew at equivalent rates when cultured separately (Figure S6C). Again, the growth rate profiles of each of these variants were unaffected by their co-culture (Figure S5D), demonstrating that no competition takes place in cultures of cells with equivalent growth rates. Conversely, culturing variant lines *v1q* and *v20q* separately or in co-culture with the faster growing variant-GFP cells showed a significantly decreased number of *v1q* and *v20q* cells within co-cultures compared to separate cultures (**Figure 3C-E**; Figure S6E,F). We confirmed that the decrease of *v1q* and *v20q* cell numbers in co-cultures with variant-GFP cells was due to apoptosis, as both sublines showed higher levels of cleaved caspase-3 staining in the co-culture condition compared to separate culture (**Figure 3F**; Figure S6G). Nonetheless, neither the decrease in cell numbers nor the level of activated caspase-3 in *v1q* and *v20q* upon competition with variant-GFP was as extensive as seen in wild-type cells upon mixing with variant-GFP. The variant lines *v20q* and *v1q* harbour an additional copy of *BCL2L1* and *MCL-1*, respectively. The higher levels of expression of anti-apoptotic proteins BCL-XL and MCL-1 (Figure S6H) are thought to confer *v20q* and *v1q* cells with increased resistance to apoptosis (Avery et al., 2013; Nguyen et al., 2014). Given that *v1q* and *v20q* variants did not assume a winner status upon mixing with wild-type cells, our data revealed that increased resistance to apoptosis is not sufficient to confer a winner cell phenotype. Together, these results confirmed that cell competition behaviour is context-dependent, and that the faster proliferation rate is a feature of variant hPSCs exhibiting a winner cell phenotype. Moreover, we show that increased resistance to apoptosis is not sufficient to confer cells with a winner cell phenotype, but it reduces the rate of loser cell elimination.

**Figure 3.**
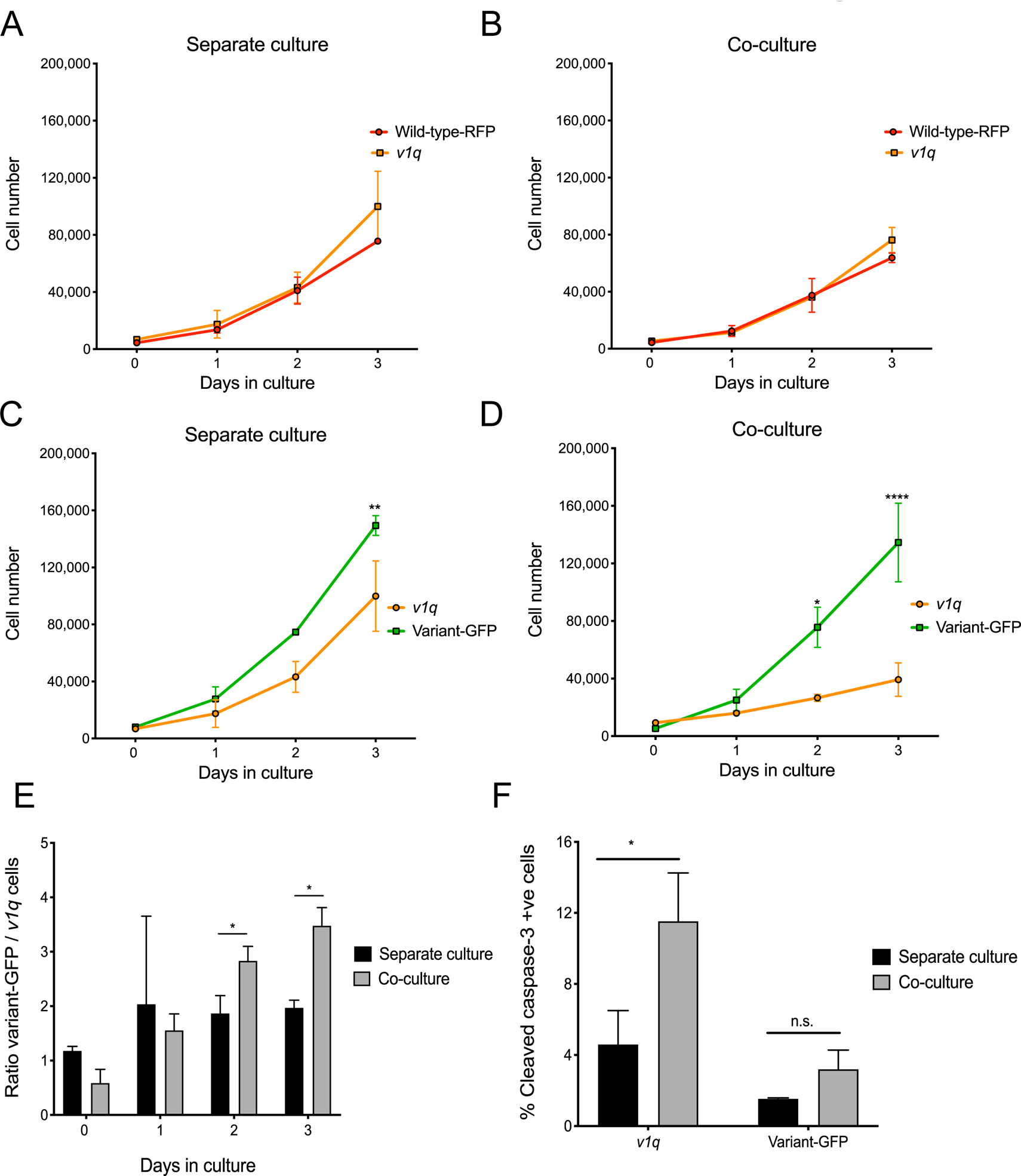
The winner phenotype is dependent on higher proliferative rates of variant cells. A) Growth curves of wild-type-RFP and *v1q* cells grown separately. B) Growth curves of wild-type-RFP and *v1q* cells grown in co-culture. C) Growth curves of *v1q* and variant-GFP cells grown separately. D) Growth curves of *v1q* and variant-GFP cells grown in co-culture. E) Ratio of variant-GFP/*v1q* cells in separate versus co-culture conditions. ‘Separate culture’ ratio was calculated by dividing the number of variant-GFP cells in separate culture with a number of *v1q* cells in separate culture. ‘Co-culture’ ratio was obtained by directly counting the number of either *v1q* or variant-GFP cells in co-culture using high-content microscopy and dividing by the total cell count. F) Percentage of cells positive for cleaved caspase-3 indicator of apoptosis in *v1q* and variant-GFP cells in separate culture or upon co-culture. A-E: Data are the mean of two independent experiments ± SD; F: Results are the mean of three independent experiments ± SD. n.s. non-significant; * p<0.05; ** p<0.01; **** p<0.0001, Student’s *t* test.

### YAP mediates the winner versus loser cell phenotype in hPSCs

To determine how cell competition in hPSC cultures is mediated at the molecular level, we initially performed transcription analysis of loser (*v1q*) and winner (variant-GFP) cells in separate and co-cultures (**Figure 4A**). We first focused on identifying expression signatures associated with prospective winner and loser populations, by analysing the differential gene expression between these cells in separate cultures (**Figure 4B,C**). In line with the complex aneuploidy of variant-GFP cells, the number of differentially expressed genes in winner versus loser cells was large, with 3524 genes significantly upregulated and 3311 genes significantly downregulated in winner compared with loser cells (**Figure 4C**). The Kyoto Encyclopedia of Genes and Genomes (KEGG) enrichment analysis (Kanehisa et al., 2010) showed that the most significantly enriched molecular network from the downregulated genes was the ribosomal pathway, followed by the cell cycle, TGF-*β* and Hippo pathway (**Figure 4D**). The Hippo signalling pathway, a key regulator of cellular fates, was also significantly enriched in the KEGG analysis of differentially expressed genes between winner and loser cells upon co-culture (**Figure 4E,F**). Given the apparent differences in the Hippo pathway between winner and loser cells, and also considering its known role in mechanical signalling (Codelia et al., 2014), we next asked whether the Hippo pathway is mediating cell competition in hPSC cultures.

**Figure 4.**
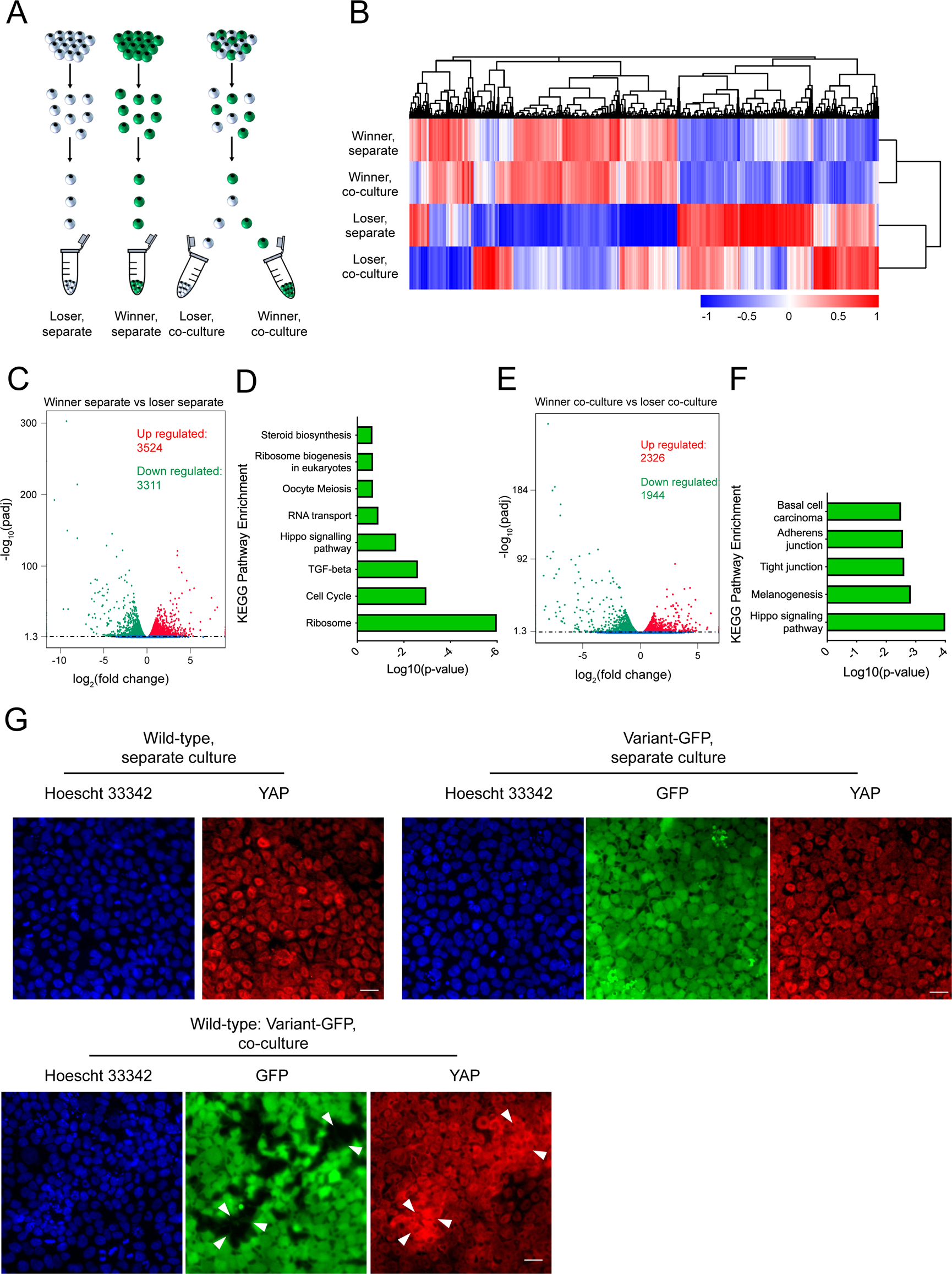
Gene expression analysis of winner and loser cells indicates Hippo signalling as a mediator of cell competition in hPSC cultures. A) Schematic depicting the sorting of loser (*v1q*) and winner (variant-GFP) cells from separate or co-culture conditions to obtain the following populations: ‘loser separate’, ‘winner separate’, ‘loser co-culture’ and ‘winner co-culture’. Four biological replicates of each sample were obtained from independent experiments. B) Unsupervised hierarchical clustering of the winner and loser cells from separate and co-culture based on the differentially expressed genes. C) Volcano plot of the differentially expressed genes between winner and loser hPSCs in separate cultures. Downregulated genes (green) are positioned on the left of the plot and upregulated genes (red) are on the right of the plot. D) KEGG pathway analysis of the downregulated genes in winner versus loser hPSCs in separate cultures showing the molecular pathways with a corrected p-value >0.25 threshold. E) Volcano plot of the differentially expressed genes between winner and loser hPSCs in co-culture. Downregulated genes (green) are positioned on the left of the plot and upregulated gene (red) are on the right of the plot. F) KEGG pathway analysis of the downregulated genes in winner versus loser hPSCs in co-culture showing the molecular pathways with a corrected p-value >0.25 threshold. G) Immunocytochemistry staining for YAP (red) in wild-type and variant-GFP cells (green) in separate cultures and upon co-culturing revealed cytoplasmic localisation of YAP in crowded wild-type cells upon co-culture. Nuclei are counterstained with Hoechst 33342. Scale bar: 25µm.

A major effector of the Hippo signalling is the transcriptional co-activator Yes-associated protein 1 (YAP), which regulates gene expression of target genes through binding to the TEA domain DNA-binding family of transcription factors (TEAD) (Zhao et al., 2008). YAP localises to the nucleus when Hippo signalling is low, whereas active Hippo signalling results in YAP phosphorylation by LATS1/2 kinase and its cytoplasmic retention (reviewed in (Totaro et al., 2018)). Notably, YAP was shown to be modulated by mechanical signalling, including mechanical stresses imposed by neighbouring cells (reviewed in (Panciera et al., 2017)). We first checked YAP localisation in separate and mosaic cultures of wild-type and variant-GFP cells by immunofluorescence. YAP localised to the nucleus of both wild-type and variant-GFP cells when they were grown in separate cultures (**Figure 4G**). Strikingly, whilst the variant-GFP cells retained the nuclear YAP in co-cultures with wild-type cells, the wild-type cells within the same culture exhibited a shift in YAP localisation from nuclear to cytoplasmic (**Figure 4G**).

To directly address the hypothesis that YAP is mediating the super-competition behaviour of hPSCs, we overexpressed YAP in wild-type cells (Figure S7A,B) and analysed the effect of overexpression on the growth and behaviour of these cells in mosaic cultures. YAP overexpression resulted in the improved growth rates and increased homeostatic density of wild-type cells, suggesting an increased threshold to mechanical sensitivity imposed by neighbouring cells (**Figure 5A**). Further, YAP overexpressing cells exhibited a winner phenotype in co-cultures with wild-type cells (**Figure 5B**; Figure S7C). Finally, in comparison with wild-type cells, YAP overexpressing cells were more resistant to crowding caused by co-culture with variant-GFP cells as evidenced by higher numbers of YAP overexpressing cells persisting in co-cultures with variant-GFP cells (**Figure 5C**; Figure S7D). Based on these results, we concluded that YAP is mediating cell competition in hPSC cultures.

**Figure 5.**
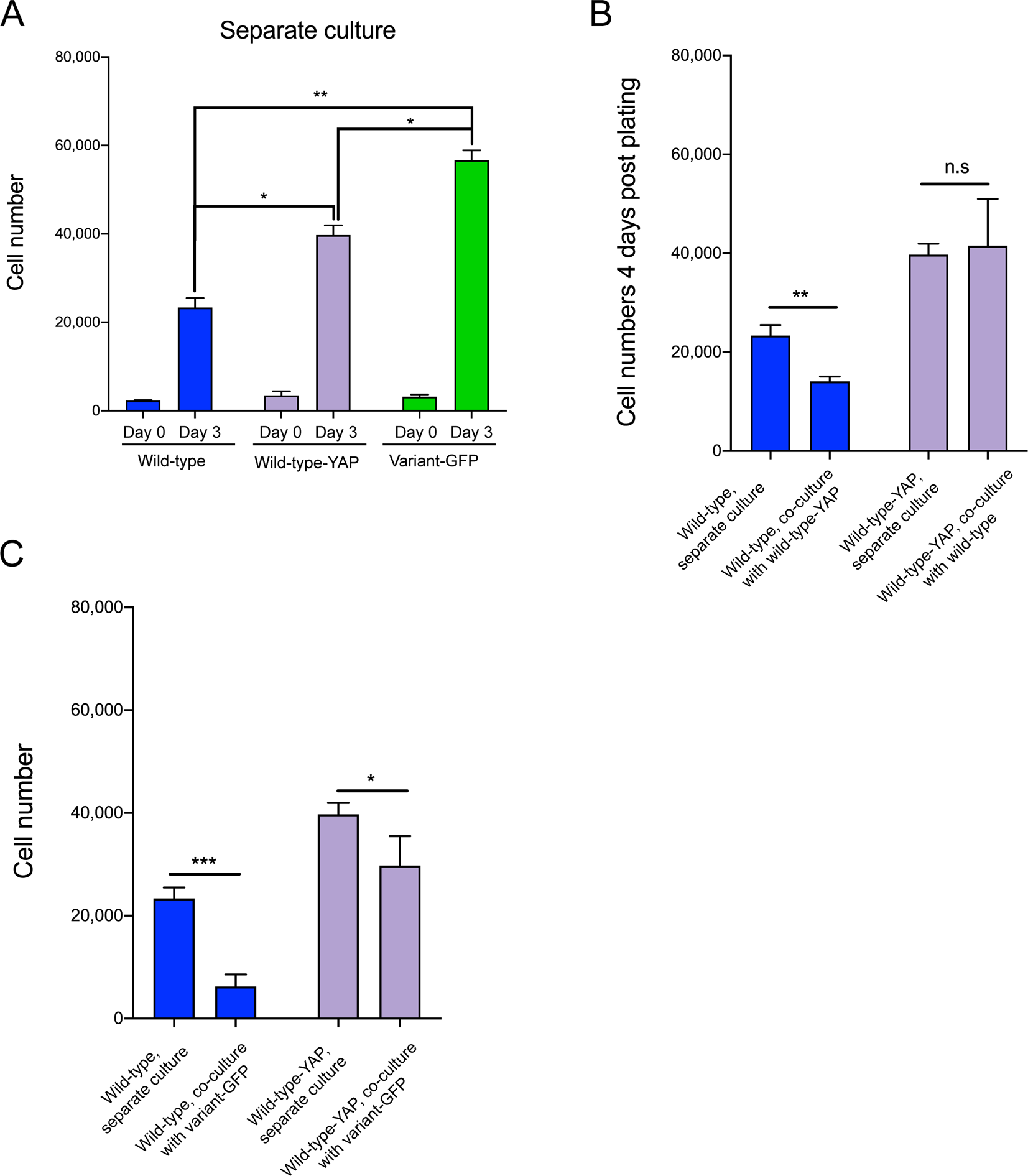
YAP overexpression alleviates the loser cell phenotype in wild-type cells. A) YAP overexpression leads to improved growth of wild-type cells. B) YAP overexpressing cells assume the winner phenotype in co-cultures with wild-type cells. C) YAP overexpression in wild-type cells confers increased resistance to cell crowding in co-cultures with variant-GFP cells. Data are the mean of three independent experiments ± SD. n.s. non-significant; *p< 0.05; **p<0.01; ***p<0.001; Student’s *t* test.

### Apical actin constriction regulates YAP localisation in hPSCs

To gain further mechanistic insight into YAP-mediated hPSC competition, we set out to investigate the upstream regulators of YAP in this context. Our observation that wild-type hPSCs are corralled into smaller spaces upon co-culture with variants, coupled with the findings from other cell models that YAP localisation can be mechanically influenced by cell shape and actin fibers (Aragona et al., 2013; Wada et al., 2011), prompted us to examine the cytoskeleton as a potential regulator of YAP in hPSCs. Phalloidin staining of F-actin showed a similar basal-to-apical profile of actin fibers in wild-type and variant-GFP cells in separate cultures, with both populations exhibiting a faint staining of actin filaments encircling the cell within the adhesion belt (Figure S8A). However, whilst the variant cells retained a similar actin distribution upon co-culture with wild-type cells, the crowded wild-type cells showed a dramatic change in their actin fibre network (**Figure 6A**). Specifically, we detected a redistribution of actin stress fibers within the adhesion belt, evident as intense staining of F-actin within the circumferential actin ring (**Figure 6A**). Expression of myosin IIB, a major non-muscle myosin, was also upregulated in the adhesion belt of the crowded wild-type cells (**Figure 6B**), reflecting the increased constriction of the adhesion belt in these cells upon co-culture with variants. The cytoplasmic YAP in wild-type cells was phosphorylated at Ser127 residue (**Figure 6C**), a known phosphorylation target of LATS1/2 kinase (Zhao et al., 2007), indicating that Hippo signalling is activated in wild-type hPSCs upon co-culture with variant cells, thus leading to sequestration of YAP in the cytoplasm of wild-type hPSCs.

**Figure 6.**
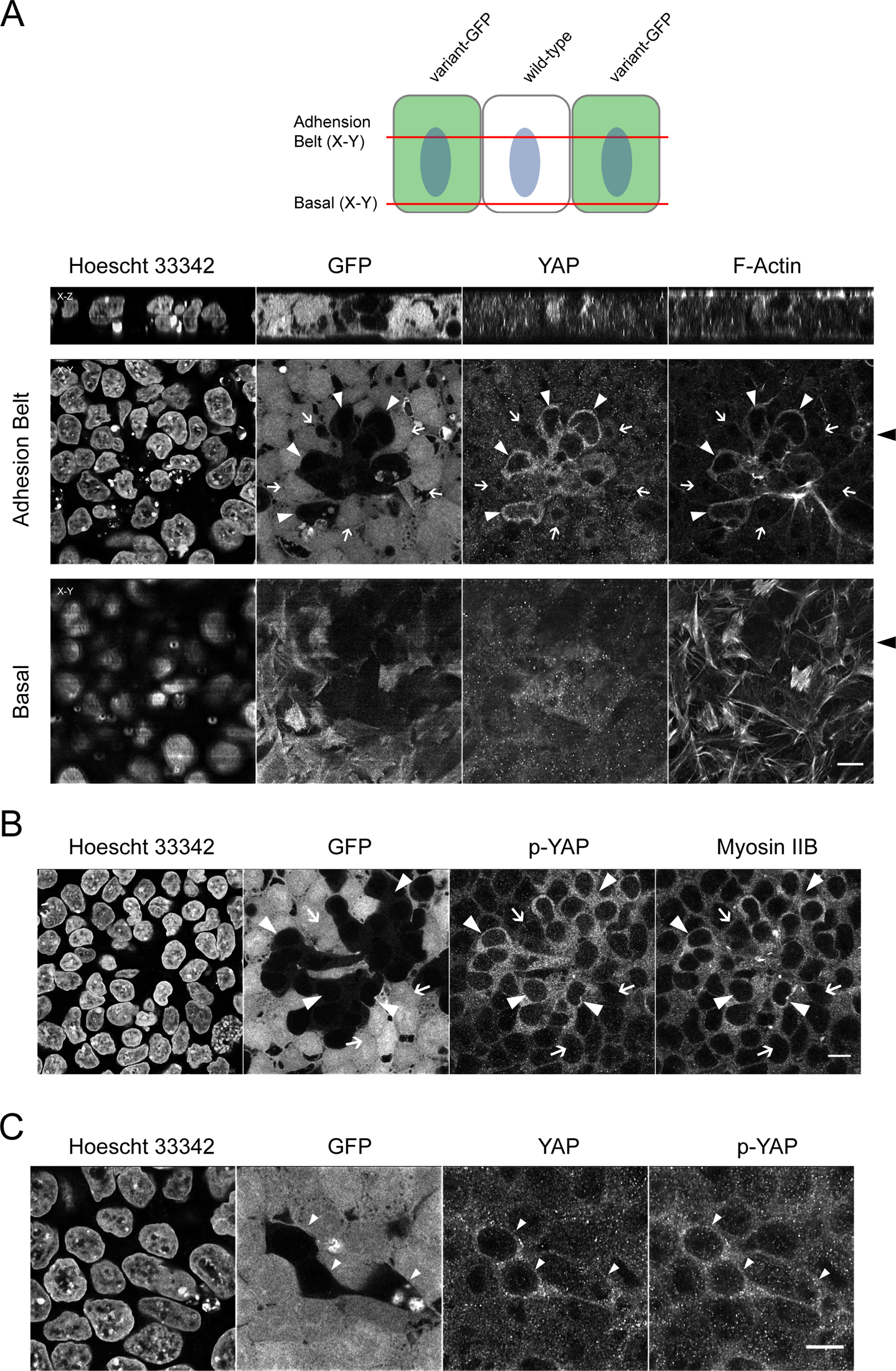
Crowded wild-type cells display prominent actin adhesion belt and cytoplasmic YAP localisation. A) Schematic representation of the adhesion belt and basal planes in confocal imaging of co-cultured cells (upper panel). Corresponding adhesion belt and basal planes of co-cultured cells stained for F-actin and YAP are shown in the panels below. Closed arrowheads point to wild-type cells displaying YAP localised within the cytoplasm and having a prominent staining of adhesion belt F-actin. Open arrows point to neighbouring variant-GFP cells displaying nuclear localisation of YAP and no prominent adhesion belt. B) Phosphorylated YAP (p-YAP) localisation and myosin IIB staining in co-cultured wild-type and variant-GFP cells taken at the plane of the adhesion belt. Closed arrowheads point to wild-type cells displaying increased p-YAP localised within the cytoplasm and having prominent staining of myosin IIB. Open arrows point to neighbouring variant-GFP cells displaying weaker phosphorylated YAP staining and less prominent myosin IIB staining. C) YAP and p-YAP staining in co-cultured wild-type and variant-GFP cells taken at the plane of the adhesion belt. Closed arrowheads point to wild-type cells displaying increased p-YAP localised in the cytoplasm. Scale bars: 10μm.

To determine whether the observed cytoskeletal differences in winner and loser cells upon co-culture underpin the differences in their sub-cellular YAP localisation, we utilised a set of chemicals that perturb actinomyosin cytoskeleton. First, we used nocodazole to disrupt microtubules. Microtubule disruption, evident by diminished *α*-tubulin staining (**Figure 7A**), reduced the adhesion belt contraction in crowded wild-type cells (**Figure 7A**). Concomitantly, we detected a shift from a predominantly cytoplasmic YAP in co-cultured wild-type cells to a diffuse (i.e. both cytoplasmic and nuclear) localisation in their nocodazole-treated counterparts (**Figure 7A**). Furthermore, YAP phosphorylation at serine 127 was suppressed upon nocodazole treatment of co-cultured wild-type cells (Figure S8C), confirming the lower levels of inactive form of YAP. Disruption of actin fibers using latrunculin A or cytochalasin B also resulted in reduced actin ring within the adhesion belt of crowded wild-type cells and a diffuse localisation of YAP in those cells (**Figure 7B**). On the other hand, inhibition of myosin activity by treating cells with the Rho-associated coiled coil kinase (ROCK) inhibitor had no overt effect on the sub-cellular localisation of YAP in wild-type and variant cells upon co-culture (**Figure 7C**). As Y-27632 changed the actin stress fibers at the cell: extracellular matrix level, but did not reduce the intense actin staining within the adhesion belt of the crowded wild-type cells (**Figure 7C**), this data suggests that a constricted adhesion belt, rather than actin stress fibers, promotes cytoplasmic localisation of YAP in hPSCs. Taken together, we conclude that in hPSC cultures super-competitive variant cells corral wild-type counterparts into areas of significantly higher density compared with the density of wild-type separate cultures. Consequent restructuring of actin fibers within the adhesion belt of crowded wild-type hPSCs causes sequestering of YAP in their cytoplasm and triggers them to commit to apoptosis.

**Figure 7.**
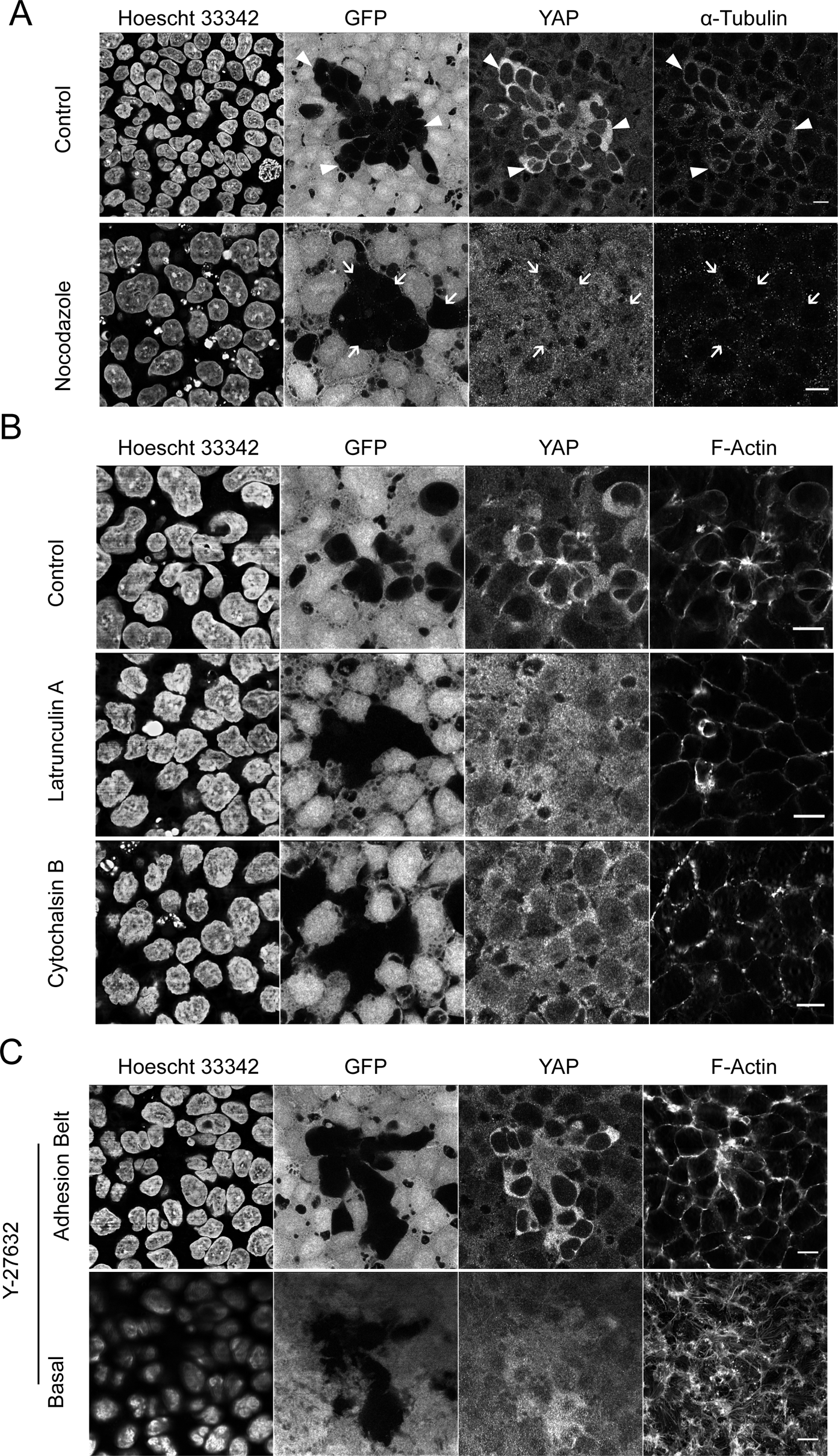
YAP localisation is regulated by adhesion belt actin in hPSCs. A) Localisation of YAP in co-cultured wild-type and variant-GFP cells treated with nocodazole. Closed arrowheads point to wild-type cells displaying YAP localised within the cytoplasm and having prominent staining of *α*-tubulin. Nocodazole treatment perturbed the microtubule structure and caused diffuse localisation of YAP in wild-type cells (open arrows). B) Disruption of F-actin in the adhesion belt of co-cultured wild-type and variant-GFP cells treated with latrunculin A or cytochalasin B resulted in the diffuse YAP localisation. C) Treatment of co-cultures with Y-27632 affected the stress fibers at the basal plane, but did not disrupt the actin in the adhesion belt. Y-27632 had no impact on the YAP localisation in the crowded wild-type cells. Scale bars: 10μm.

## Discussion

Suppressing the commonly arising variant hPSCs from overtaking the cultures requires a thorough understanding of the attributes that facilitate variant cells in achieving the clonal dominance. Here we report that the supremacy of particular variant clones in hPSC cultures is enhanced through competitive interactions with their wild-type counterparts, leading to the elimination of wild-type cells from mosaic cultures. The manner of wild-type cell elimination resembles previously described cell competition (Mamada et al., 2015; Morata and Ripoll, 1975; Sancho et al., 2013) in that the wild-type hPSCs, albeit viable in homotypic cultures, failed to thrive and underwent increased levels of apoptosis when co-cultured with variants. We showed that the competitive behaviour in hPSC context was not mediated by soluble factors, as the co-culture conditions in which winner and loser cells shared the same media, but had no direct cell contacts, did not cause apoptosis of loser cells. Instead, the winner cell phenotype was assumed by variant clones which possessed relatively faster growth rates and achieved higher homeostatic density compared to the loser cells. Thus, cell competition in hPSC cultures is akin to mechanical cell competition, which is characterised by faster-growing winner cells causing compaction and subsequent elimination of the slower-growing losers (Levayer et al., 2016; Wagstaff et al., 2016).

Given that variant hPSCs in homotypic cultures displayed a lower propensity to apoptosis compared with wild-type cells, differential sensitivity to apoptosis could be a plausible explanation for the selective elimination of loser cells upon competition for space with the variant counterparts. However, two lines of evidence from our study suggest that the resistance to mechanical forces, rather than apoptosis *per se*, determines the winner versus loser status in the context of mosaic hPSC cultures. First, variant lines which possessed increased resistance to apoptosis, but did not exhibit a proliferative advantage and increased homeostatic density compared to wild-type cells, did not display a winner phenotype upon mixing with wild-type counterparts. Conversely, we detected a cell competition phenotype only upon mixing two sublines with differential proliferation rates, with a relatively faster subline adopting a winner status. Secondly, our observation that neighbouring loser and winner cells display differential distribution of mechanosensitive transcription regulator YAP, despite coexisting in the same culture, suggested that winner and loser hPSCs interpret their mechanical environments differently.

The remarkable changes in the shape of loser cells when corralled by variants, and the finding that variants displace the wild-type cells in cell confrontation assays, indicated a differential sensitivity of wild-type and variant cell populations to crowding upon competition for space in hPSC cultures. Whether the crowding sensing involves sensing cell volume and shape or direct sensing of mechanical forces (Valon and Levayer, 2019) remains unknown, nonetheless, our data point to the actomyosin cytoskeleton as a key mediator of crowding sensing in hPSCs. Indeed, we detected significant changes in the cytoskeleton of crowded loser cells, which displayed prominent staining of actin fibers within the adhesion belt. Accordingly, we showed that disruption of F-actin by cytochalasin B or latrunculin A, decouples the crowd sensing from YAP localisation, as YAP was retained in the nucleus of crowded loser cells treated with actin inhibitors. Although F-actin regulation of YAP has been previously noted (Aragona et al., 2013; Dupont et al., 2011), disruption of F-actin primarily led to cytoplasmic and not, as we observed, the nuclear localisation of YAP. The stark contrast of our findings with those previously published fits with the notion that the actomyosin activity has differing effects on the localisation of YAP at low and high cell density. This difference has been attributed to the remodelling of the actin cytoskeleton from predominantly stress fibers to an apical actin ring at low and high density, respectively (Furukawa et al., 2017). Analogous to our findings, the actin inhibition in high cell density MDCK cells displaying a prominent circumferential belt and cytoplasmic YAP localisation, promoted re-distribution of YAP to nucleus (Furukawa et al., 2017). Therefore, it follows that the effect of actin cytoskeleton on YAP localisation is context and possibly cell type-dependent. Further investigation is warranted in order to delineate the exact mechanism by which hPSCs integrate environmental cues through their actin cytoskeleton to cease expansion, and how this mechanism may be evaded by variant cells in mechanical cell competition.

YAP was previously implicated in mechanical cell competition of several different experimental systems. In cells of aggressive brain tumors, glioblastomas, the level of YAP expression determines the winner versus loser cell status and contributes tumorigenesis by promoting the expansion of a clone with a higher expression of YAP and increased expression of downstream tumorigenic genes (Liu et al., 2019). In NIH3T3 embryonic fibroblasts, the transcription factor TEAD and its regulator YAP, have been also found to control cell proliferation and competition (Mamada et al., 2015). Perhaps of most significance for hPSC biology, TEAD-YAP axis was also shown to control cell competition in the pluripotent cells of the mouse epiblast (Hashimoto and Sasaki, 2019). In this context, cell competition was found to be a quality control mechanism regulating elimination of unspecified cells within the embryo. However, in contrast to various cell types grown *in vitro,* in pre-implantation mouse embryos YAP localisation appears decoupled from growth inhibition (Nishioka et al., 2009). Whether such decoupling holds true in post-implantation embryos and to what extent the sensitivity to crowding of hPSCs reflects biology of their *in vivo* counterparts is currently unknown.

The ability of some of the commonly acquired variant hPSCs to tolerate higher cell densities is reminiscent of transformed cells which evade the contact inhibition to achieve hyperproliferation. In numerous cancer cells, YAP is often either overexpressed or activated (Zanconato et al., 2016), although mutations in YAP itself, or additional genes within the Hippo pathway, are altogether relatively rare (Harvey et al., 2013). Given that commonly amplified regions in hPSC genome typically span several megabases (Baker et al., 2016), it is difficult to pinpoint potential driver from mere passenger mutations implicated in culture adaptation of hPSCs. Nonetheless, neither *YAP* nor several other key genes of the Hippo pathway (e.g. *WWTR1*, *TEAD1*, *LATS1*, *LATS2* and *NF2*) map to the chromosomes commonly amplified in variant hPSCs. Hence, based on the data thus far and drawing on parallels with the cancer field, it is tempting to speculate that pathways upstream of Hippo, may be affected by genetic changes in hPSCs. Based on the data in our study, genes implicated in the regulation of actin cytoskeleton and cell proliferation would be the prime candidates. In that respect, it is worth noting the recent identification of recurrent point mutations in hPSCs, which entail mutations in genes implicated in cytoskeleton and control of the cells cycle (Avior et al., 2019). It will be interesting to evaluate the behaviour of such mutants in the context of super-competition to narrow down the candidate genes that underpin this phenotype.

Whilst identification of driver genes in hPSC variants and their relationship to YAP regulation awaits further analyses, the observation that the variants exert their advantage through mechanical cell competition suggests that spatial constraints may have provided an important selective pressure leading to selection and fixation of commonly occurring variants. This notion becomes particularly likely when viewed in the light of the early reports of hPSC cultures that advocated hPSCs to be grown at a high cell density, due to the need for cell-cell contacts in sustaining hPSC proliferation and survival (Fox et al., 2008; Thomson et al., 1998). Thus, it stands to reason that high cell density conditions created an environment for mechanical cell competition due to the lack of available space in expanding hPSC cultures. Intriguingly, not all commonly occurring variants that we tested displayed a super-competitive advantage in hPSC cultures, likely reflecting adaptation to different types of selective pressures, other than cell crowding. Ultimately, detailed characterisation of variants will allow not only the identification of conditions that select for them, but importantly, will also enable stratification of genetic variants as a necessary requirement for risk assessment of cellular therapy products. At least in theory, the most concerning would be the genetic variants that impinge on the behaviour of surrounding non-variant cells in a manner as described in this study.

Together, our results point to a model whereby mechanical cues from hPSC environment dictate the stem cell fate. The findings of our study hold a number of implications for the use of hPSCs in research and regenerative medicine. First, the finding that actin cytoskeleton mediates YAP localisation and ultimately cell fate in hPSCs opens up opportunity to harness this knowledge in order to control hPSC behaviour. Secondly, our conclusion that winner/loser status in hPSC cultures is determined by relative proliferative abilities of cells and YAP localisation provides a potential indicator that could be tested in order to allow stratification of potentially detrimental variants in the context of regenerative medicine. Finally, our results showing that cell crowding mediates survival advantage of the variants, coupled with the finding that high cell density conditions promote genome damage due to the media acidification (Jacobs et al., 2016), suggest that scaling up of hPSC cultures must be executed at carefully controlled cell densities.

In conclusion, our work revealed cell competition as an important aspect of cellular interaction of wild-type and variant hPSCs, contributing to the genetic drift that culminates in a complete overtake of cultures by super-competitive variant clones. Undertaking further detailed analyses of genetic variants that exhibit super-competitive behaviour should be informative for impact on regenerative medicine applications.

## METHODS

### Human pluripotent stem cell (hPSC) lines

Wild-type hPSCs used in this study were early passage sublines of H7 (WA07) and H14 (WA14), originally established in the laboratory of James Thomson (Thomson et al., 1998), which were karyotypically normal (based on at least 20 metaphases analysed by G-banding of cell banks prior to experiments and at various time points upon subsequent passaging) and did not possess a commonly gained 20q11.21 copy number variant (as determined by quantitative PCR for copy number changes and/or Fluorescent In Situ Hybridisation (Baker et al., 2016)). Spontaneous variants with karyotypic abnormalities were detected during the subsequent culture of H7 and H14 cells at the Centre for Stem Cell Biology in Sheffield (Baker et al., 2007; Draper et al., 2004). Genetically variant sublines of H7 line used in this study and their karyotypes were: ‘variant-GFP’ cells [48,XX,+del(1)(p22p22),der(6)t(6;17)(q27;q1),+12)] (30 metaphases analysed), also harbouring chromosome 20q CNV as determined by quantitative PCR analysis and FISH (Baker et al., 2016); ‘*v1,17q,i20*’ [47,XX, +del(1)(p22p22), der(6)t(6;17)(q27;q1), t(12;20)(q13;q11.2), i(20)(q10) dup(20)(q11.21q11.21)] (30 metaphases analysed); and ‘*v1q*’ cells [46,XX,dup(1)(q21q42)] (30 metaphases analysed). The variant ‘*v20q*’ appeared to have a diploid karyotype when analysed by G-banding (30 metaphases analysed), but a gain of a copy number variant 20q11.21 was detected by Fluoresecent In Situ Hybridisation and quantitative PCR analysis. The karyotype of the H14 variant subline H14.BJ1-GFP was 48,XY,+12,+der(17)hsr(17)(p11.2) del(17)(p13.3) (20 metaphases analysed). Variants *v1q* and *v20q* were established in this study by cloning out spontaneously arising variants from mosaic cultures using single cell deposition by fluorescent activated cell sorting. Single cells from mosaic cultures were sorted directly into individual wells of a 96 well plate using a BD FACS Jazz and cultured to form colonies over 2-3 weeks. The resulting colonies were expanded in culture and subsequently frozen to establish cell banks. At the time of freezing, sister flasks were sent for karyotyping by G-banding and assessment of the relative copy number of commonly identified genetic changes by qPCR as described below.

### Human pluripotent stem cell (hPSC) culture

Flasks used for hPSC maintenance were coated with vitronectin (VTN-N) (Cat. # A14700, Life Technologies) diluted to 5 µg/ml in Dulbecco’s phosphate buffered saline (PBS) and incubated at 37°C for 1h prior to aspirating the vitronectin solution and plating hPSCs. HPSCs were maintained in E8 medium prepared in house, consisting of DMEM/F12 (Cat. # D6421; Sigma-Aldrich) supplemented with 14 µg/l sodium selenium (Cat. # S5261; Sigma-Aldrich), 19.4 mg/l insulin (Cat. # A11382IJ; Thermo Fisher Scientific), 543 mg/l NaHCO_3_ (Cat. # S5761; Sigma-Aldrich), 10.7 mg/l transferrin (Cat. # T0665; Sigma-Aldrich), 10 ml/l Glutamax (Cat. # 35050038; Thermo Fisher Scientific), 100µg/l FGF2 (Cat. # 100-18B; Peprotech) and 2 µg/l TGF*β*1 (Cat. # 100-21; Peprotech) (Chen et al., 2011). For time lapse experiments, E8 was prepared using DMEM/F12 without phenol red (Cat. # D6434; Sigma-Aldrich). Cells were fed daily and maintained at 37°C under a humidified atmosphere of 5% CO_2_ in air. Routine passaging every 4-5 days was performed using ReLeSR (Cat. # 05873; STEMCELL Technologies) according to manufacturer’s instructions. Cells were resuspended in E8 and split at 1:3 or 1:4 ratio (wild type cells) or 1:8 to 1:30 ratio (variant sublines). Cells were genotyped after thawing and every 5-8 passages by G-banding, Fluorescent In Situ Hybridization and/or using quantitative PCR for common genetic changes.

### Karyotyping by G-banding

Karyotyping by G-banding was performed by the Sheffield Diagnostic Genetics Service (https://www.sheffieldchildrens.nhs.uk/sdgs/). To capture the cells in metaphase, hPSC cultures were treated with 0.1 µg/ml KaryoMAX Colcemid Solution in PBS (Cat. # 15212012; Life Technologies) for 2 - 4h. Cells were then harvested with 0.25% trypsin/versene (Gibco, Invitrogen) and pellets re-suspended in pre-warmed 0.0375M KCl hypotonic solution. Following a 10 min incubation in KCl, cells were pelleted again and fixed with methanol:acetic acid (3:1). Metaphase spreads were prepared on glass microscope slides and trypsin solution briefly spread over the slides prior to staining with 4:1 Gurr’s/Leishmann’s stain (Cat. # L6254; Sigma-Aldrich). Slides were scanned, images of banded metaphases captured and analysed using the Leica Biosystems Cytovision Image Analysis system (version 7.5 build 72). At least 20 metaphases were analysed.

### Fluorescent In Situ Hybridisation (FISH) for 20q copy number variant

FISH for chromosome 20q copy number variant was performed by the Sheffield Diagnostic Genetics Service (https://www.sheffieldchildrens.nhs.uk/sdgs/). Cells were harvested and pelleted at 270 *g* for 8 min. The cell pellet was resuspended in 0.0375 M potassium chloride pre-warmed to 37°C. After a 10 min incubation at room temperature, the cells were fixed in methanol:acetic acid (3:1). A small volume (∼50µl) of cell suspension was dropped onto glass slides. The interphase FISH was performed using the *BCL2L1* probe covering the genes *BCL2L1*, *COX4I2* and 3’ end of *ID1* (green fluorescently labelled BAC (RP5-857M17) provided by BlueGnome (Illumina)) and the 20q telomere probe the TelVysion 20q Spectrum Orange (Cat. # 08L52-001; Abbott). The cells on slides and probes were denatured by heating up to 72°C for 2 min in a PTC-200 DNA Engine (Peltier Thermal Cycler, MJ Research). Hybridisation was performed at 37°C for 16h. Slides were washed in 0.4x sodium citrate with 0.3% Tween 20 and 2x sodium citrate with 0.1% Tween 20. Coverslips were mounted on the slides in 20µl, Vectashield Mounting Medium with DAPI (**Cat.** #: H-1200; Vector Laboratories). One hundred interphase cells were analysed on an Olympus BX51 fluorescent microscope.

### Quantitative PCR (qPCR) for determining copy number changes of target genes

Relative copy number of commonly identified genetic changes was assessed using the qPCR-based approach described in (Baker et al., 2016). Genomic DNA was extracted from hPSCs using the DNeasy Blood & Tissue Kit (Cat. # 69504; QIAGEN) and digested with FastDigest EcoRI (Cat. # FD0275; Thermo Fisher Scientific) for 2 h at 37°C, followed by inactivation at 65°C for 20 min. PCR reactions were set up in triplicate, with each 10µl PCR reaction containing 1X TaqMan Fast Universal Master Mix (Cat. # 4352042; Thermo Fisher Scientific), 100nM of forward and reverse primers (**Table S1**), 100nm of probe from the Universal Probe Library (**Table S1**) and 10ng of genomic DNA. PCR reactions were run on a QuantStudio 12K Flex Thermocycler (Cat. # 4471087; Life Technologies). Following the first two steps of heating the samples to 50°C for 2 min and denaturing them at 95°C for 10 min, reactions were subjected to 40 cycles of 95°C for 15 s and 60°C for 1 min. The Cq values were obtained from the QuantStudio 12K Flex Software with auto baseline settings and were then exported to Excel for copy number analysis using the relative quantification method (2^-ddcq^). The calibrator samples for the qPCR assay were hPSC gDNA samples previously established as diploid using karyotyping and Fluorescent In Situ Hybridisation analyses (Baker et al., 2016).

### Cell competition assay

Cells were dissociated to single cells using TrypLE (Cat. # 11528856; Thermo Fisher Scientific) for 4 min at 37°C, washed once in DMEM/F12, counted and resuspended in E8 media supplemented with 10µM Y-27632 (Cat. # A11001-10; Generon). Cells were plated as separate cultures of each subline or mixed cultures of different sublines, as described in the individual experiments. After 24h, the medium was removed and the wells were washed once with basal medium DMEM/F12 (Cat. # D6421; Sigma-Aldrich) to remove the Y-27632. The medium was replaced with E8 and that point was considered as ‘day 0’ of competition experiments. Cells were cultured for further 72 hours and fed daily with E8 medium. Cells were fixed at different time points post-plating in 4% paraformaldehyde (PFA) for 15 min at room temperature, and nuclei stained with 10µg/ml Hoechst 33342 (Thermo Fisher Scientific). In every mixing experiment, one of the sublines used was fluorescently labelled (e.g. either variant-GFP mixed with other non-labelled sublines or wild type-RFP mixed with other wild type or variant sublines), thus allowing identification of cell numbers of each of the sublines in mixed cultures. Imaging of the entire 96 well was performed using the InCell Analyzer (GE Healthcare) high-content microscopy platform. Quantification of total and individual subline cell numbers was performed either using custom protocols in Developer Toolbox 1.7 software (GE Healthcare) or CellProfiler (Carpenter et al., 2006).

For growth curve analysis, cells were plated at 4,4×10^4^ cells/cm^2^ in separate cultures or co-cultures, with the co-cultures containing 50:50 ratio of different sublines (i.e. 2,2 x10^4^ cells/cm^2^ of each subline). As an additional control, separate cultures were also plated containing equivalent numbers of cells from co-cultures (i.e. 2,2 x10^4^ cells/cm^2^ for each subline). Cells were fixed with 4% PFA at different time points post-plating and the cell numbers analysed as described above.

For assessing the effect of increasing ratios of variant cells on wild-type cell growth, wild type and variant-GFP cells were plated in E8 supplemented with 10µM Y-27632 (Cat. # A11001-10; Generon) at the total number of 4,4×10^4^ cells/cm^2^, with the ratio of variant cells varying from 10% to 90% of the total cell number. After the initial 24h post-plating, cells were washed with DMEM/F12 (Cat. # D6421; Sigma-Aldrich) to remove the Y-27632 and then grown in E8 for further 3 days. Cells were then fixed with 4% PFA and the cell numbers analysed as described above.

For assessing the effect of increasing cell density on wild-type cell growth, wild type and variant-GFP cells were plated at a 50:50 ratio, at cell densities increasing from 3,750 to 45,000 cells/cm^2^. After the initial 24h post-plating, cells were washed with DMEM/F12 (Cat. # D6421; Sigma-Aldrich) to remove the Y-27632 and then grown in E8 for further 3 days. Four days post-plating, cells were fixed with 4% PFA and the cell numbers analysed as described above.

### Time-lapse imaging and analysis

Time-lapse microscopy was performed at 37°C and 5% CO_2_ using a Nikon Biostation CT. Cells were imaged every 10 min for 72 h using 10x or 20x air objective. Image stacks were compiled in CL Quant (Nikon) and exported to FIJI (Image J) (Schindelin et al., 2012) for analysis. Lineage trees were constructed manually from FIJI movies. Individual cells were identified in the first frame and then tracked in each subsequent frame until their death, division or the end of the movie. The timing of cell death or division for each cell was noted and then used to reconstruct lineage trees of founder cells using either TreeGraph 2 (Stover and Muller, 2010) or Interactive Tree Of Life (iTOL) (Letunic and Bork, 2007) software.

### Transwell assay

For indirect co-culture, Millipore Transwell 8.0μm PET membrane inserts (Cat. # PIEP12R48; Millipore) were used in combination with 24 well plates. Both the insert and well were coated with vitronectin (VTN-N) (Cat. # A14700, Life Technologies) diluted to 5 µg/ml in PBS. Cells were harvested using TrypLE (Cat. # 11528856; Thermo Fisher Scientific) and 1.5×10^4^ cells were seeded in the well and insert. Cells were pre-cultured independently for 24h in E8 medium supplemented with 10μM Y-27632 (Cat. # A11001-10; Generon) to facilitate cell attachment. 24h post-plating, cells were washed with DMEM/F12 (Cat. # D6421; Sigma-Aldrich) to remove the Y-27632 and inserts were subsequently placed into appropriate wells with fresh E8 medium. Medium was changed daily until the end of the experiment when the cells were fixed with 4% PFA.

### Cell confrontation assay

Cells were harvested using TrypLE (Cat. # 11528856; Thermo Fisher Scientific) and washed once in DMEM/F12 (Cat. # D6421; Sigma-Aldrich). After counting, 5×10^4^ cells were seeded in E8 medium supplemented with 10µM Y-27632 (Cat. # A11001-10; Generon) into the inner compartment of two-well silicone inserts (Ibidi 80209). One day post-plating the silicone inserts were removed, leaving a defined 500μm gap between the two cell populations. The cells were then washed with DMEM/F12 (Cat. # D6421; Sigma-Aldrich) to remove Y-27632 and the medium was replaced with fresh E8 medium. Cells were fed daily and left to grow for four days until the two opposing cell fronts had been in contact for approximately 48h. Cells were then fixed with 4% PFA for 15 min at room temperature, followed by washing in PBS. Cells were subsequently stained for the apoptotic marker cleaved caspase-3 (Cat. #:9661; Cell Signaling Technology) and nuclei were counterstained with Hoechst 33342 (Cat. # H3570; Thermo Fisher Scientific). Images were processed in CellProfiler (Carpenter et al., 2006) to identify wild-type, wild-type-RFP and variant-GFP cells. Using the nuclei stain, each cell was assigned a positional identity relative to the border and further analyzed for positive cleaved caspase-3 signal. Using the positional information of each cell, figures displaying the location of each cell, as well as cleaved caspase-3 positive cells were constructed in R (R Project for Statistical Computing; RRID:SCR_001905).

### Local density analysis

To compute the local density of each cell, the data was processed in the programming language R. Delaunay triangulation was performed on each image by using the cell nuclei as points for the triangulation. For each cell, the sum of areas of Delaunay triangles sharing a vertex with the cell of interest was calculated. As this sum is inversely proportional to the compactness of the cells, local cellular density is taken as the inverse of this sum. Mathematically, the local density ρ for each cell is defined as:

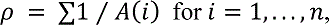

Where n is the number of Delaunay triangles that share a vertex with the cell of interest, and A(i) is the area of Delaunay triangle i.

### Immunocytochemistry

Cells were fixed with 4% PFA for 15 min at room temperature, and permeabilised with either 0.5% Triton-X in Dulbecco’s phosphate buffered saline (PBS) for 10 min or 0.2% Triton-X in PBS for 1h. Cells were then incubated with 1% bovine serum albumin (BSA) and 0.3% Triton X-100 in PBS. Primary and secondary antibodies, their suppliers and the dilutions used are listed in the Key Resources Table. Cells were incubated with primary antibodies either for 1h at room temperature or overnight at 4°C with gentle agitation on an orbital shaker. Following three washes with PBS, cells were incubated with an appropriate secondary antibody in PBS supplemented with 1% BSA, 0.3% Triton X-100 and 10µg/ml Hoechst 33342 for 1h at 4°C. Cells were then washed three times with PBS before imaging. Cells that were prepared for confocal imaging were grown on glass coverslips and mounted onto slides in 20 µl Vectashield Mounting Medium (**Cat.** #: H-1000; Vector Laboratories). Images were captured using the InCell Analyzer (GE Healthcare) or ZEISS LSM 880 (Carl Zeiss AG, Oberkochen, Germany) fitted with an Airyscan detection unit.

### Flow cytometry

Flow cytometry for cleaved caspase-3 was performed to assess levels of apoptotic cells in cultures. To collect apoptotic cells which had detached from the flask, the old media was added to a 5ml FACS tube and centrifuged at 270 x *g* for 5 min. Remaining cells in the flask were harvested with TrypLE (Cat. # 11528856; Thermo Fisher Scientific) and added to the FACS tube containing the collected cells from the supernatants of the same flasks. The collated sample was pelleted and the cell pellet fixed in 4% PFA for 15 min at room temperature. Cells were permeabilised with 0.5% Triton X-100 in PBS for 5 min at room temperature and then incubated with anti-cleaved caspase-3 primary antibody (Cat. # 9661; Cell Signalling Technology) in the blocking buffer (1% BSA and 0.3% Triton X-100 in PBS). Samples were gently agitated for 1h at room temperature, prior to washing three times in blocking buffer and staining with secondary antibody (Goat anti-Rabbit AffiniPure IgG+IgM (H+L), Cat. # 111-605-003-JIR; Stratech) for 1h at room temperature in the dark. Cells were then washed twice with blocking buffer and analysed on BD FACS Jazz. Baseline fluorescence was set using secondary antibody-only stained samples.

For intracellular analysis of YAP, cells were harvested with TrypLE, permeabilised and blocked as described above. Cells were incubated with anti-YAP antibody (Cat. # sc-101199; Santa Cruz Biotechnology) and gently agitated for 1h at room temperature, prior to washing three times in blocking buffer and staining with secondary antibody (Goat anti-Mouse AffiniPure IgG+IgM (H+L), Cat. # 115-605-044-JIR; Stratech) for 1h at room temperature in the dark. Cells were then washed twice with blocking buffer and analysed on BD FACS Jazz. Baseline fluorescence was set using secondary antibody-only stained samples.

### Cell sorting of individual sublines from co-cultures

After establishing that variant *v1q* cells are losers in co-cultures with variant-GFP cells, we performed mixing experiments of *v1q* and variant-GFP cells in T75 flasks, following the same protocol as in 96 well plates. Briefly, cells were plated at 4,4×10^4^ cells/cm^2^ in E8 supplemented with Y-27632 (Cat. # A11001-10; Generon). After 24 h, Y-27632 was removed and cells were cultured in separate or mixed cultures for another day. Cells were then harvested using TrypLE (Cat. # 11528856; Thermo Fisher Scientific) for 4 min at 37°C, washed with DMEM/F12 (Cat. # D6421; Sigma-Aldrich), counted and resuspended at 2×10^6^ cells/ml in E8 media. Sorting was performed using BD FACSJazz cell sorter (BD Biosciences). Sort gates were set using the separate culture unlabelled *v1q* cells and variant-GFP separate cultures as baseline and positive gates, respectively. GFP-negative *v1q* and GFP-positive variant-GFP cells were sorted into collection vessels at 5×10^5^ cells per sample. Samples were re-analysed post sorting to establish the purities. In all cases a minimum purity of 98% was achieved. Separate cultures were also put through the same sorting procedure as co-cultures. Samples were centrifuged at 270 x *g* for 3 min, supernatant removed and cell pellets stored at −80°C prior to RNA or protein extraction. Samples from four independent experiments were obtained for further analyses.

### RNA extraction, sequencing and bioinformatic analysis

Four biological replicates of *v1q* and variant-GFP cells FACS-sorted from either separate or mixed cultures were used for RNA extraction and RNAseq analysis. The RNA was isolated using a Qiagen RNAeasy Plus Mini Kit (Cat. # 74134; Qiagen), and the RNA concentration and purity determined using a Qubit 3.0 Fluorometer (Life Technologies, Carlsbad, USA) and NanoPhotometer (Implen, Munich, Germany), respectively. The libraries were constructed and sequenced by Novogene (Beijing, China). Briefly, libraries were prepared using NEBNext Ultra RNA Library Prep Kit for Illumina (New England Biolabs, Ipswich, USA) and the library preparations were sequenced on an Illumina Hiseq platform (Illumina, San Diego, USA) to generate 150 bp paired-end reads. The sequencing reads were aligned to a reference human genome using TopHat v2.0.12. Raw read counts were calculated using the HTSeq v0.6.1 and were normalized into the fragments per kilobase of transcript per million mapped reads (FPKM), based on the length of the gene and reads count mapped to it. Differential gene expression analysis was performed using the DESeq R package (1.18.0). Genes with the Benjamini and Hochberg’s adjusted p value of < 0.05 were considered differentially expressed. To identify potential signaling pathways within differentially expressed genes, KEGG enrichment analysis of differentially expressed genes was performed using the PANTHER v14 software (Mi et al., 2019). The resulting list was refined using REViGO (Supek et al., 2011) to remove redundant GO terms.

### Western blotting

Cells were lysed in 1x Laemmli Buffer pre-warmed to 95°C and the total protein concentration was normalised using the Pierce BCA Protein Assay (Cat. # 23250; ThermoFisher Scientific). Proteins (10µg/sample) were resolved by SDS-PAGE and were run alongside a Page Ruler prestained protein ladder (Cat. # 26616; ThermoFisher Scientific). Proteins were then transferred onto a PVDF membrane (Cat. # IPVH00010; Millipore) using an Electrophoresis Transfer Cell (Bio-Rad). The membrane was blocked in 5% milk for one hour, washed three times with TBS-T (50 mM Tris-HCl (pH 7.5), 150 mM NaCl, 0.1% (v/v) Tween 20) and then incubated with primary antibodies for MCL-1 (Cat. # 5453; Cell Signalling Technology) at 1: 1,000 dilution, BCL-XL (Cat. # 2764; Cell Signalling Technology) at 1:1,000 dilution, BCL2 (Cat. # 2870; Cell Signalling Technology) at 1: 1,000 or *β*-ACTIN (Cat. #66009-1-Ig; Proteintech) at 1:5,000 dilution. Following three washes with TBS-T, the membrane was incubated with secondary antibody (either Anti-Rabbit IgG (H+L), HRP conjugate Cat. # W4011; Promega at 1:4,000 dilution or Anti-Mouse IgG (H+L), HRP conjugate Cat. # W4021; Promega at 1: 4,000 dilution) for 1h. After three washes, immunoreactivity was visualised using ECL Prime detection kit (Cat. # RPN2232, GE Healthcare) and signal captured on a CCD-based camera (Syngene).

### YAP overexpression

The pCAG-YAP expression vector was established by inserting a YAP-T2A-mCherry sequence into the pCAGeGFP vector (Liew et al., 2007). In brief, pGAMA-YAP, a gift from Miguel Ramalho-Santos (Cat. # 74942; Addgene) (Qin et al., 2016), was obtained from Addgene. Single digests were performed on the pGAMA-YAP and pCAGeGFP vectors using EcoRI (Cat. # 0101, New England Biolabs) and NotI (Cat. # 0189, New England Biolabs) restriction sites, respectively, to linearize plasmids. The cohesive ends were blunted using T4 DNA polymerase (Cat. # M0203, New England Biolabs) and vectors subsequently digested at the NheI restriction site to produce single cohesive ends. The YAP-T2A-mCherry sequence was obtained by gel extraction (Cat. # 740609, Machery-Nagel) and inserted into the pCAGeGFP using ligation reaction (Cat. # M0202, New England Biolabs) to produce the pCAG-YAP expression vector. To generate the wild-type YAP overexpressing line, cells were transfected using the Neon Transfection System (Cat. # MPK10025; Thermo Fisher Scientific). Wild-type H7 cells were dissociated to single cells using TrypLE as described above and resuspended at 2,0 x10^4^ cells/ml in “R buffer”. Transfection was performed with 5µg of plasmid DNA using 1 pulse of 1600V, 20msec width. After electroporation, the cells were immediately transferred to a vitronectin coated 60mm diameter culture dish (Cat. # 150288; Thermo Fisher Scientific) containing E8 media supplemented with 10µM Y-27632 (Cat. # A11001-10; Generon). To select for stably transfected cells, 48h post transfection cells were subjected to puromycin (Cat. # A11138; Thermo Fisher Scientific) drug selection. Individual colonies of resistant cells appeared after 1-2 weeks and were handpicked by micropipette, and transferred into a 12-well culture plate. The cells were then expanded in the presence of puromycin selection and subsequently frozen to establish cell banks. At the time of freezing, sister flasks were sent for karyotyping by G-banding and assessment of the relative copy number of commonly identified genetic changes by qPCR, as described above. Upon defrosting and subsequent culture, cells were also regularly genotyped by karyotyping and screened for common genetic changes by quantitative PCR, as described above.

### Generation of wild-type-RFP cell line

To generate the wild-type-RFP line, karyotypically diploid H7 subline was transfected with pCAG-H2B-RFP plasmid (a kind gift from Dr Jie Na, Tsinghua University, Beijing) using the 4D nucleofector (Lonza) in the “P3 Primary Cell solution” as per the manufacturer’s instructions. Cells were pulsed using the CB-150 pulse code, optimised for hPSCs. Cells were then plated into flasks coated with Geltrex (Cat. # A1413202; ThermoFisher Scientific) in mTESR1 medium (Cat. # 85850; STEMCELL Technologies) supplemented with 10µM Y-27632 (Cat. # A11001-10; Generon). After two days, the stably transfected cells were selected by selection with puromycin (Cat. # A11138; Thermo Fisher Scientific). Resistant colonies were manually picked and expanded. Clonal lines were then screened for their RFP expression levels by fluorescent imaging. The chosen clone was karyotyped by G-banding and screened for common genetic changes by quantitative PCR prior to freezing and at regular intervals (∼5 passages) upon subsequent culture.

### Treatment of hPSCs with cytoskeletal inhibitors

HPSCs were treated with either 10 µM nocodazole (Cat. #487928; VWR International), or 10 µM Y-27632 (Cat. # A11001-10; Generon) for 3h or 0.5 µM latrunculin A (Cat. # 10010630-25ug-CAY; Cambridge Bioscience) or 0.5 µM cytochalasin B (Cat. # C2743-200UL; Sigma-Aldrich) for 1h. DMSO was used as vehicle control for nocodazole, cytochalasin B and Y-27632, whereas ethanol was used as vehicle control for latrunculin B. Cells were fixed with 4% PFA for 15 min at room temperature, washed in PBS and processed for immunocytochemistry as detailed above.

## QUANTIFICATION AND STATISTICAL ANALYSIS

Statistical analysis of the data presented was performed using GraphPad Prism version 7.00, GraphPad Software, La Jolla California USA, www.graphpad.com. Differences were tested by statistical tests including Student’s *t* test or one-way ANOVA, as indicated in figure legends.

## SUPPLEMENTAL INFORMATION

Supplemental Information includes 8 figures and 4 videos.

## AUTHOR CONTRIBUTIONS

Conceived and designed the experiments: IB, TAR, CJP. Performed the experiments: CJP, IB, DS, JL. Analyzed the data: CJP, DS, PJG, IB, SS. Wrote the paper: IB, CJP, PJG, TAR.

## ACKNOWLEDGMENTS

We thank Sheffield Genetics Diagnostics Service for karyotype and Fluorescent In Situ Hybridisation analyses. This work was supported by the Medical Research Council MR/N009371/1 and the UK Regenerative Medicine Platform, MRC reference MR/R015724/1.

---

**Supplementary Figure S1.**
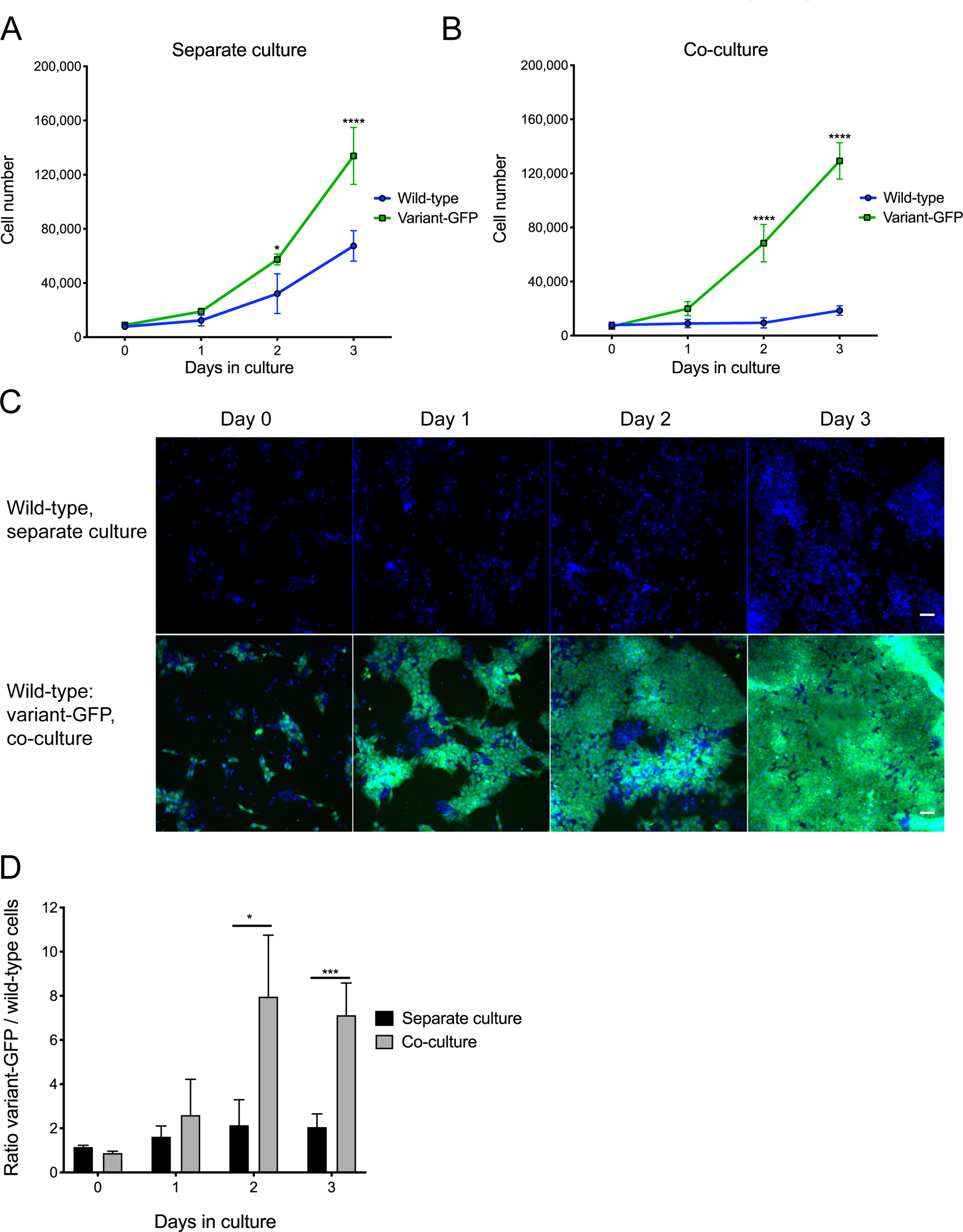

**Supplementary Figure S2.**
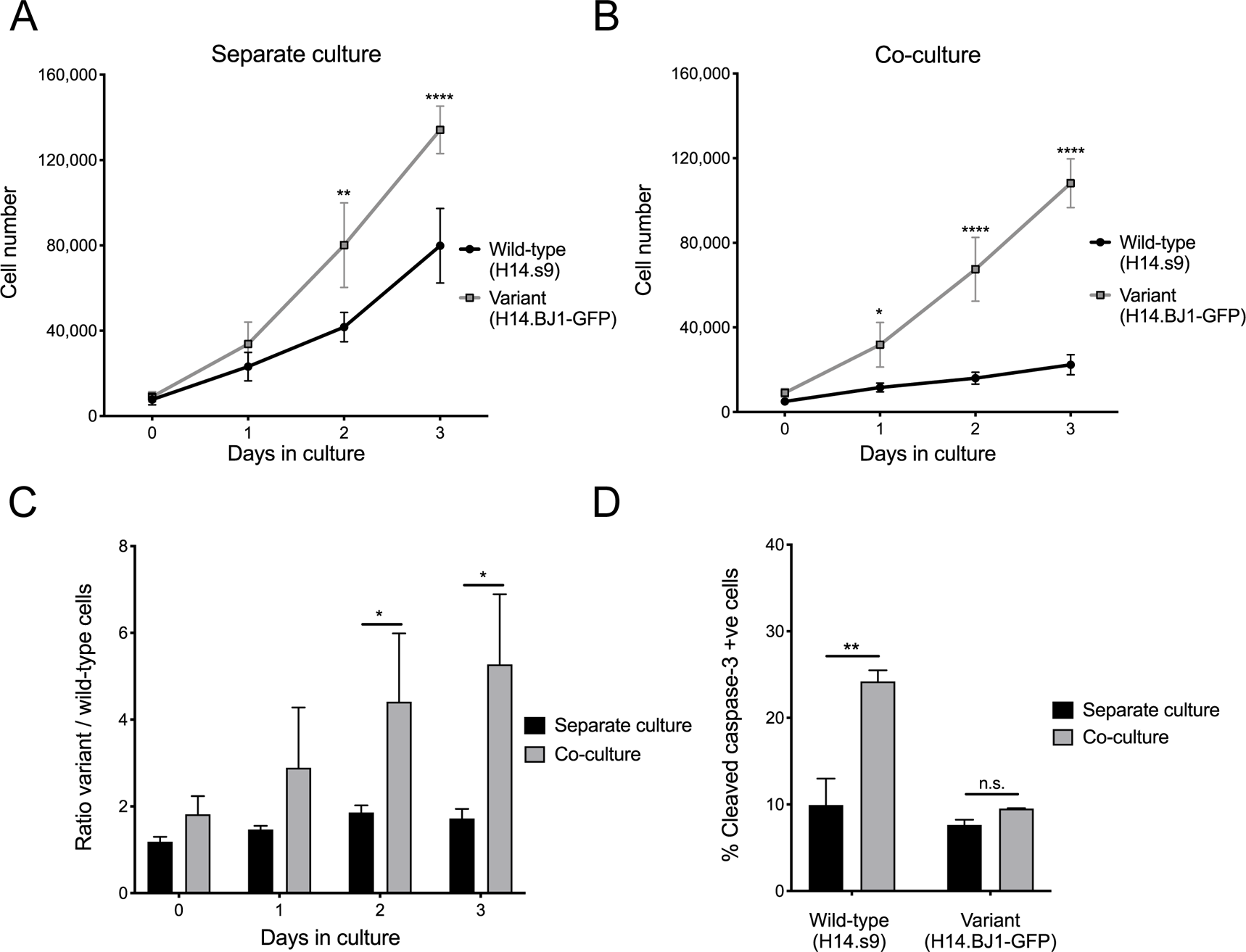

**Supplementary Figure S3.**
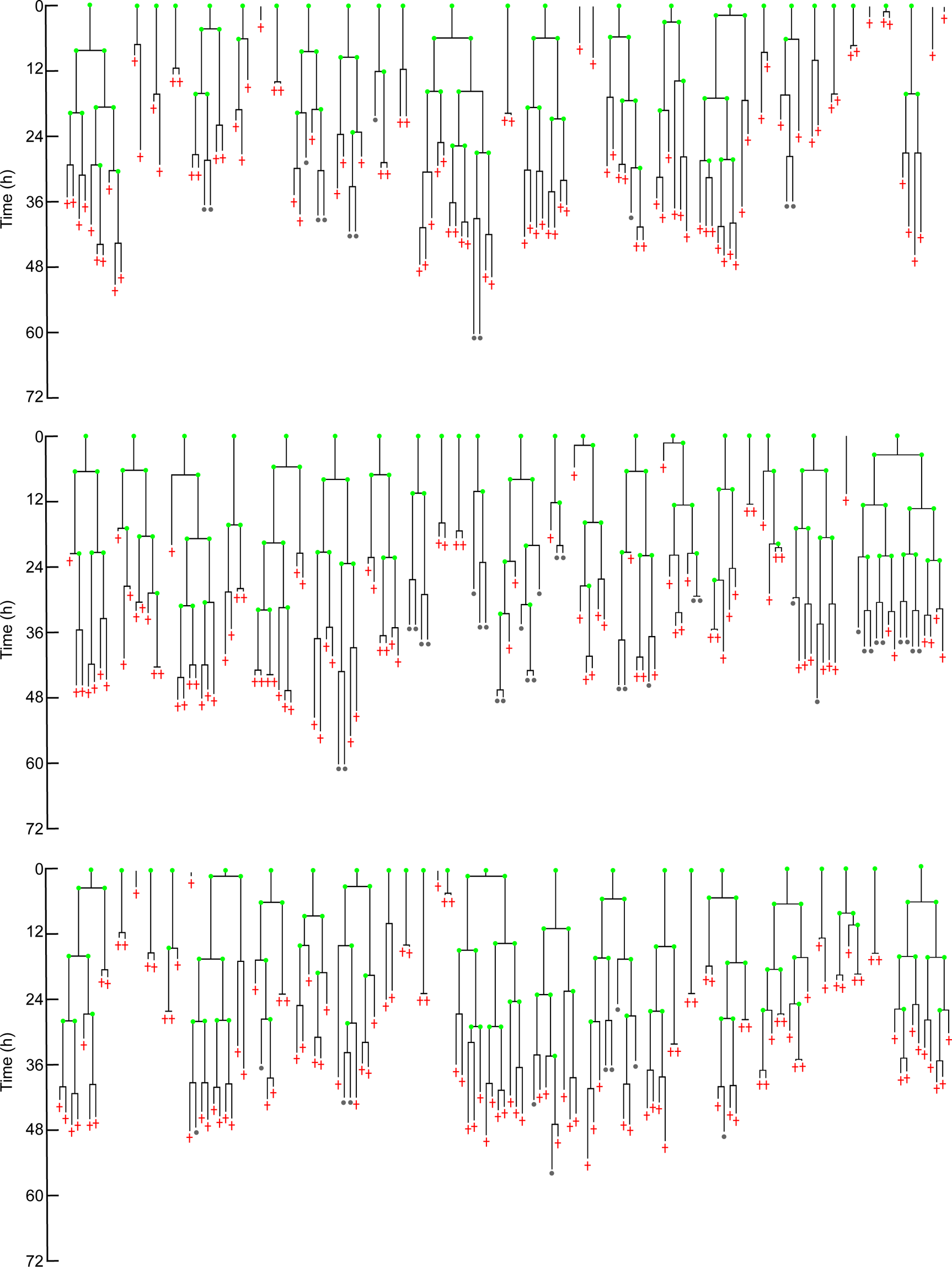

**Supplementary Figure S4.**
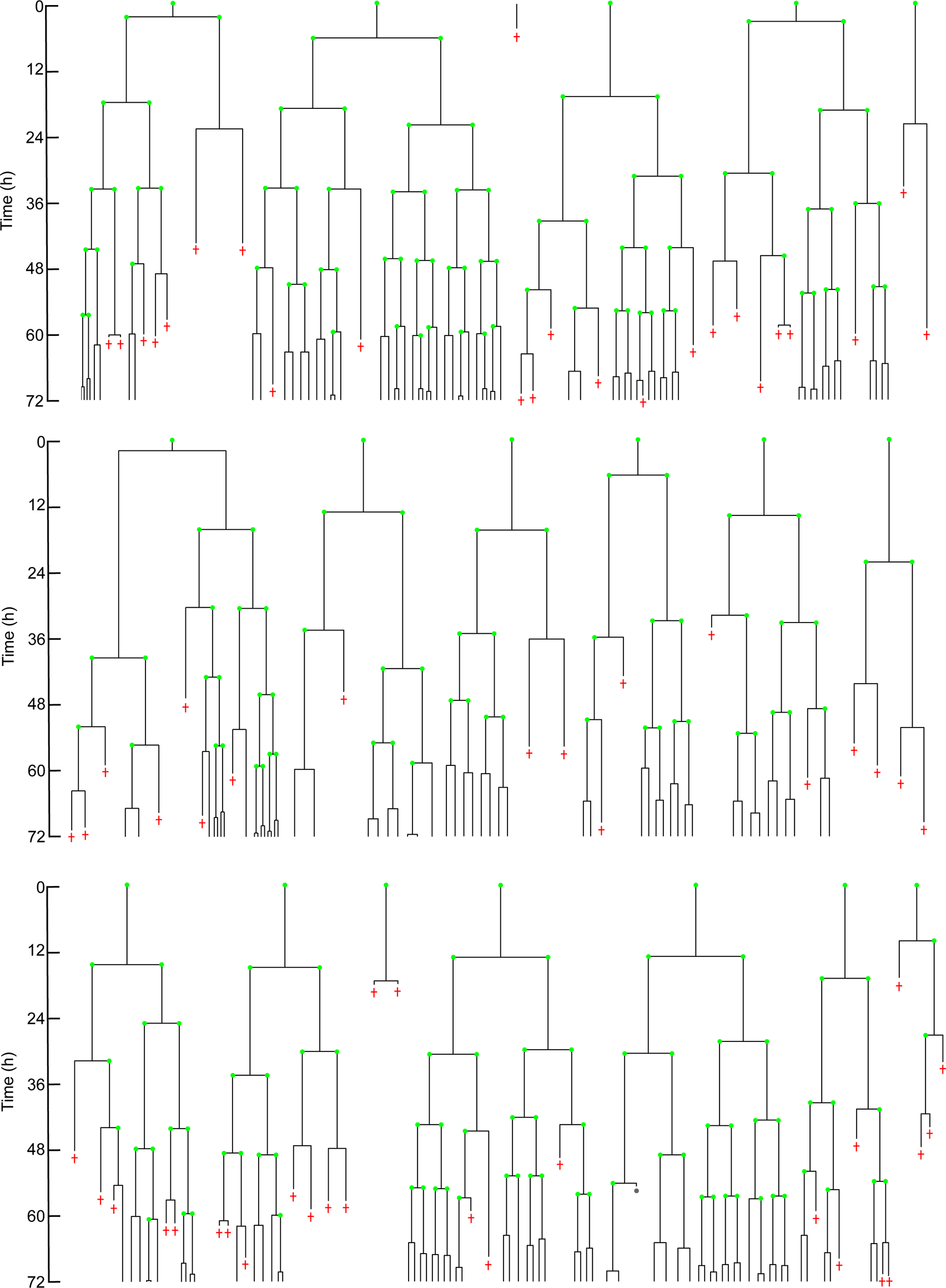

**Supplementary Figure S5.**
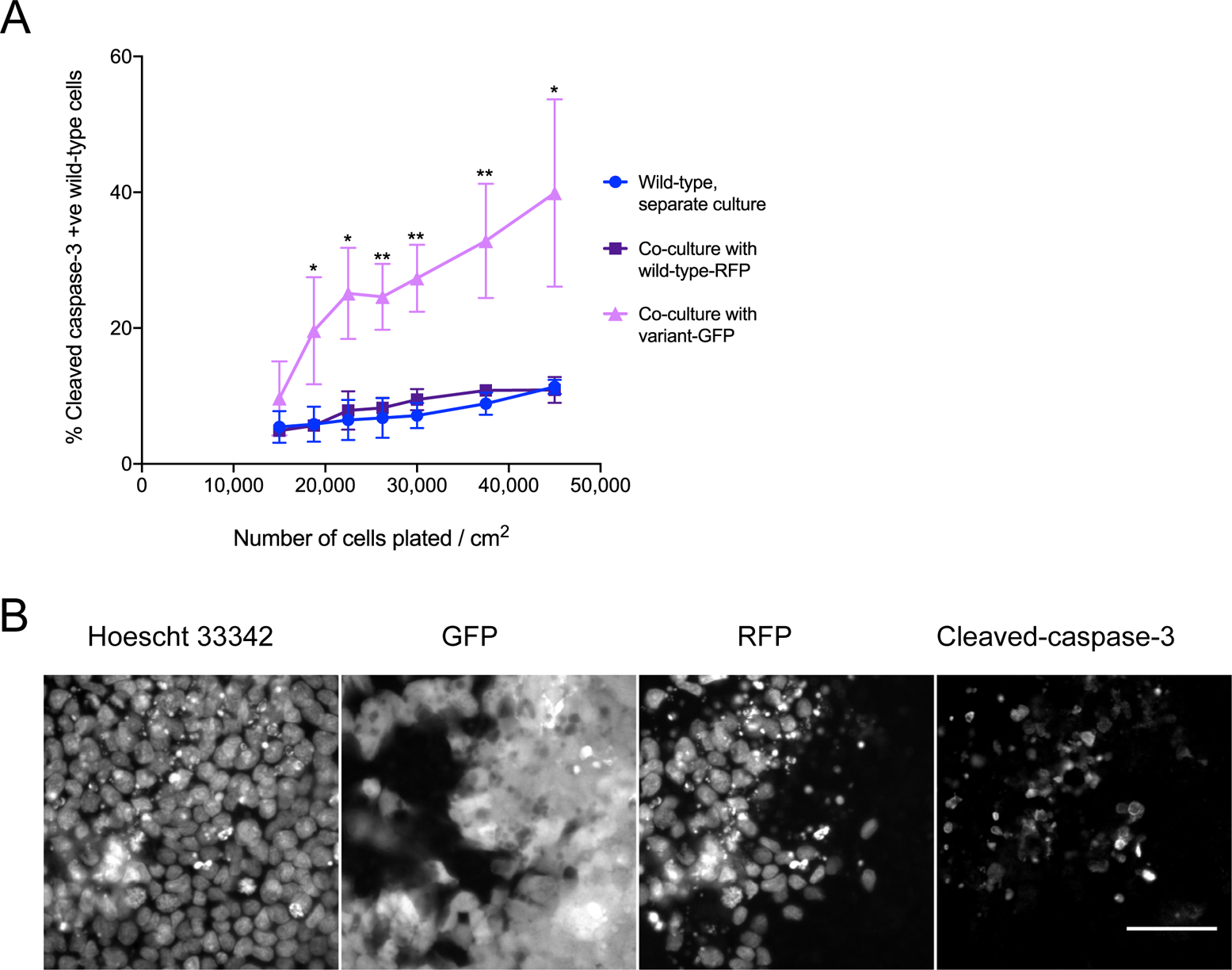

**Supplementary Figure S6.**
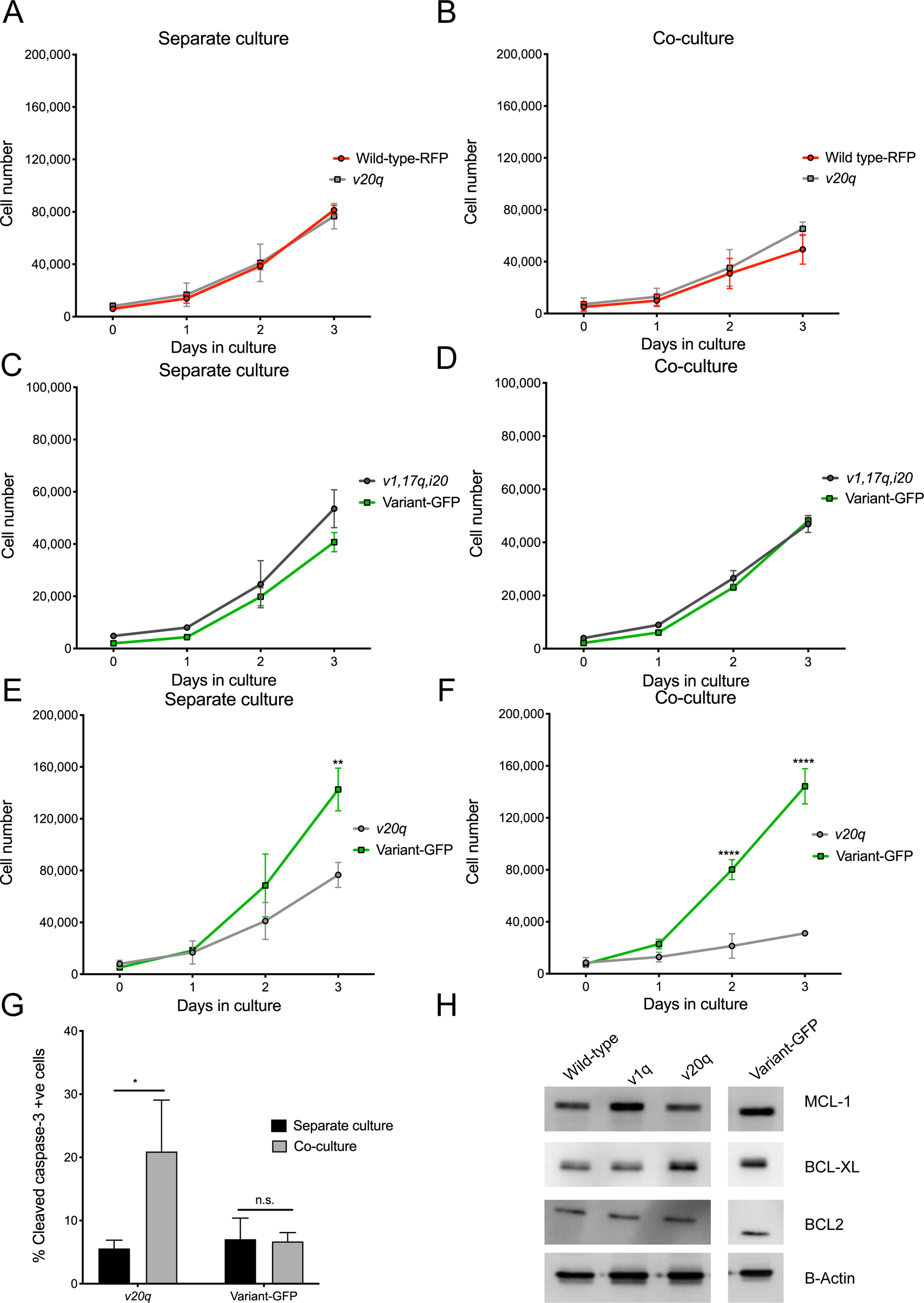

**Supplementary Figure S7.**
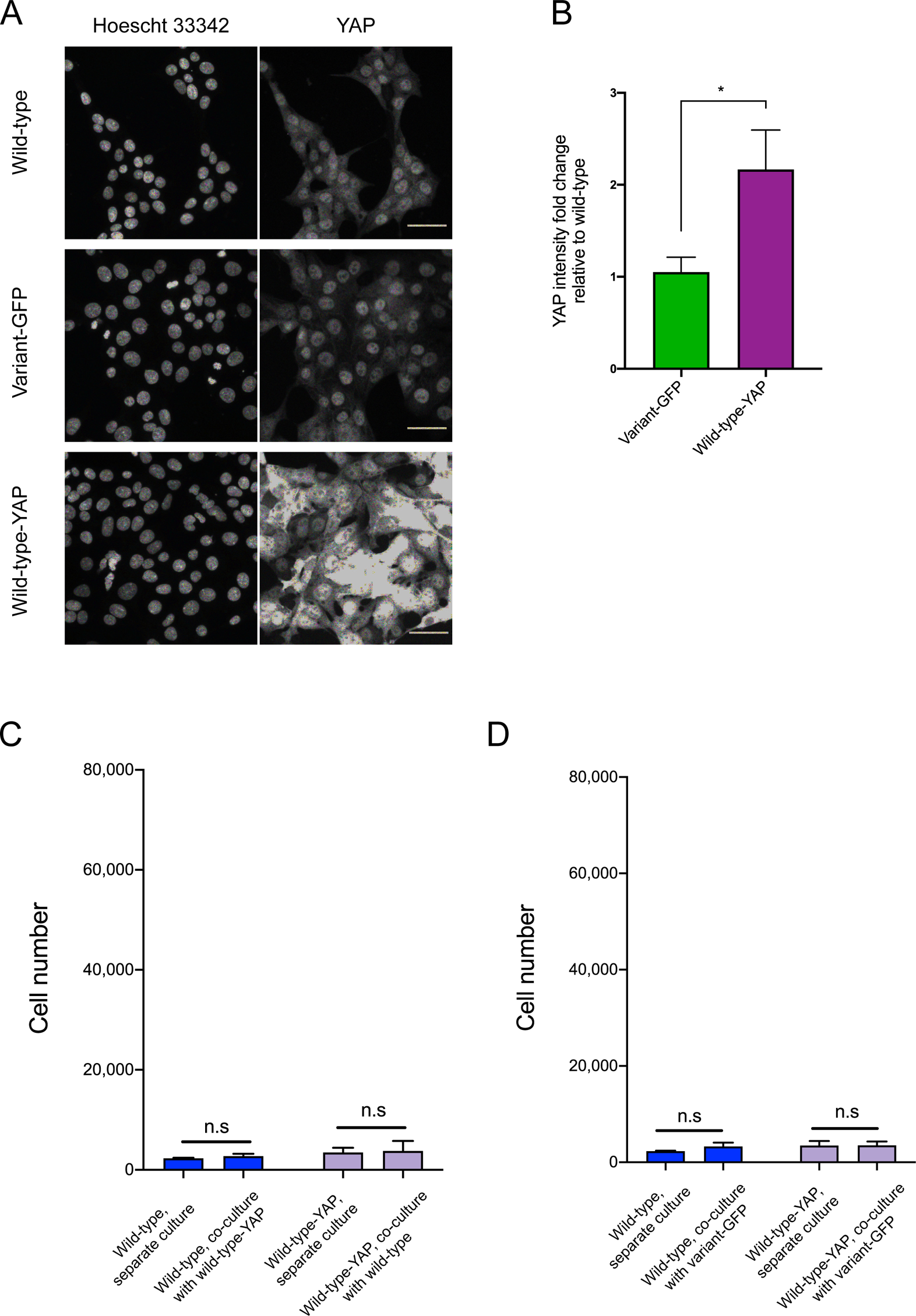

**Supplementary Figure S8.**
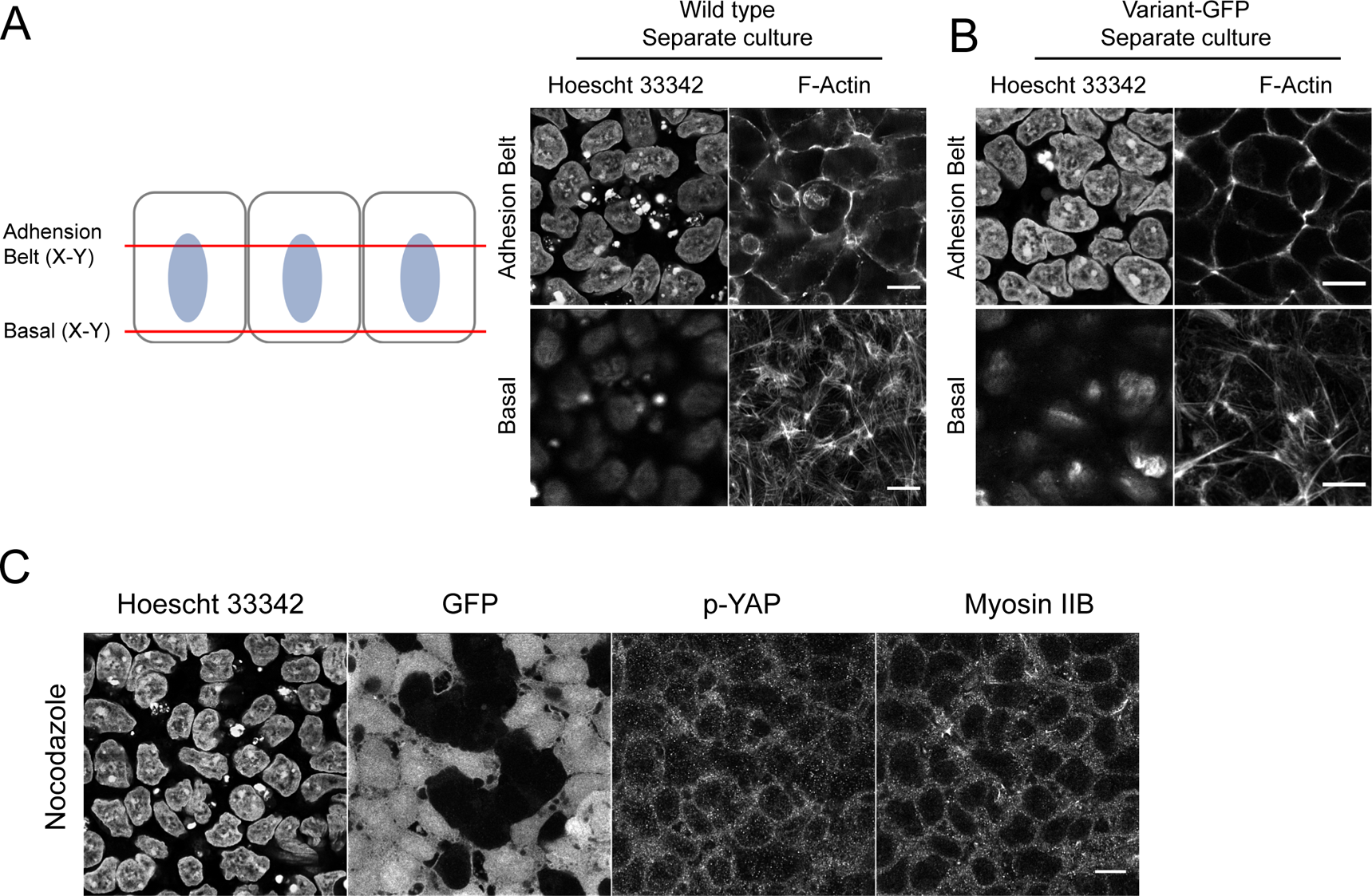

